# Uncovering the Multifaceted Role of the Kinetochore Scaffold 1 (KNL1) in Hepatocellular Carcinoma: Integrative Bioinformatics Analysis and CRISPR-Cas9 Experimental Study

**DOI:** 10.64898/2026.10.31.763338

**Authors:** Nermin M. Mohamed, Rania Hassan Mohamed, Mahmoud M. Elhefnawi, Nadia M. Hamdy

## Abstract

Hepatocellular carcinoma (HCC) is a highly aggressive malignancy characterized by pronounced heterogeneity, chromosomal instability, and limited therapeutic options. Kinetochore scaffold 1 (KNL1), a core mitotic checkpoint component, is implicated in multiple cancers, but its role in HCC remains unclear. Here, we integrated bulk and single-cell transcriptomic analyses with functional in vitro validation to elucidate the biological and clinical significance of KNL1 in HCC. KNL1 was markedly overexpressed in HCC tumors, particularly those harbouring TP53 mutations, and correlated with increased tumor break load, suggesting a link to structural chromosomal instability. Weighted gene co-expression network analysis (WGCNA) and hallmark gene set enrichment identified a KNL1-associated G2M checkpoint network, in which KNL1 ranked among the top 10 hub genes. A KNL1-G2M network score, stratified patients into high- and low-score groups, with high scores correlates with poorer overall survival, increased proliferation, and higher chromosomal instability. Immune deconvolution (CIBERSORTx) revealed that high-score tumors were enriched in T-regulatory cells and M0 macrophages, and had elevated TIDE and exclusion scores, indicating potential resistance to immune checkpoint blockade. Single-cell RNA-seq confirmed preferential activation of the KNL1-G2M module in malignant hepatocytes and highlighted increased extracellular matrix remodeling and immunomodulatory interactions. CRISPR-Cas9–mediated KNL1 downregulation in HepG2 cells induced modest G2/M arrest, suppressed proliferation, reduced survivin and Bcl2 expression, and produced ultrastructural changes consistent with early apoptosis. Collectively, our results implicate KNL1 and its associated G2/M network in tumor progression–related processes, including chromosomal instability, proliferative activity, and immune modulation in HCC. While further mechanistic and clinical studies are required, KNL1 and its associated network may represent a biologically relevant vulnerability and a potential biomarker of aggressive disease behavior in hepatocellular carcinoma.

## 1. Introduction

### 1.1. Background

Liver cancer is the third leading cause of cancer-related mortality worldwide, with hepatocellular carcinoma (HCC) accounting for approximately 75%–85% of cases (1,2). Its incidence continues to rise, driven largely by chronic viral hepatitis, excessive alcohol consumption, and metabolic syndrome (3,4). The therapeutic landscape for HCC is profoundly influenced by disease stage and liver function, with systemic therapies forming the cornerstone of management in advanced disease (5). Among these, targeted agents such as the multi-kinase inhibitors sorafenib, lenvatinib, regorafenib, cabozantinib, and ramucirumab have modestly extended patient survival; however, their benefits are often curtailed by the frequent development of drug resistance (6–11). In recent years, immune checkpoint inhibitor (ICI)-based combinations have reshaped the treatment paradigm, with regimens such as atezolizumab plus bevacizumab (anti-programmed death-ligand 1 (PD-L1) and anti-vascular endothelial growth factor (VEGF)), tremelimumab plus durvalumab (anti-cytotoxic T-lymphocyte–associated protein 4 (CTLA-4) and anti-PD-L1), and nivolumab (anti programmed cell death protein 1 (PD-1)) demonstrating promising efficacy in clinical trials (12–14). Despite these therapeutic advances, resistance, high recurrence rates, and poor long-term survival remain major challenges, highlighting the urgent need for novel treatment strategies (9,15,16). Treatment failure is driven by multiple factors, including late-stage diagnosis, primary or acquired resistance, and-at the molecular level-the pronounced intratumoral heterogeneity and complex tumor microenvironment (TME) that characterize HCC (17–20).

A critical component of this TME complexity is extracellular matrix (ECM) remodelling, which not only facilitates tumor invasion and metastasis but also contributes to immune exclusion by creating physical and biochemical barriers that impair effective anti-tumor immune responses (21,22). This dynamic remodelling of the ECM plays a pivotal role in shaping tumor progression and therapeutic resistance, highlighting the importance of deciphering the molecular drivers behind these changes in HCC (22,23). Underpinning this complexity is the profound heterogeneity of HCC, which spans multiple dimensions and significantly influences treatment response and patient prognosis (19,24–27).

Among the key drivers of HCC heterogeneity is chromosomal instability which stands out as an early and frequent event in hepatocarcinogenesis (28–30). By generating subclones with divergent karyotypes, driver mutations, and pathway dysregulation, chromosomal instability fuels both genetic and phenotypic diversity (30). It encompasses a spectrum of genomic alterations, including single nucleotide variants, somatic copy number alterations, and structural variants (31–33). Chromosomal instability can be quantified through multiple genomic metrics, including tumor mutational burden, fraction of genome altered, aneuploidy score and tumor break load, each reflecting distinct yet complementary aspects of genomic disruption (34–37).

One major contributor to chromosomal instability is mitotic dysfunction, particularly defects in kinetochore–microtubule attachment and spindle assembly checkpoint signalling, both critical for accurate chromosome segregation (38). Kinetochore Scaffold 1 (KNL1), also known as Cancer Susceptibility Candidate 5 (CASC5), is a core spindle assembly checkpoint component essential for recruiting checkpoint proteins and stabilizing kinetochore–microtubule interactions during mitosis (39).

Pan-cancer analyses highlights KNL1 as a potent oncogenic driver, with overexpression linked to poor survival in at least twelve malignancies (40). Its upregulation is frequent in pancreatic, hepatocellular carcinomas, lung, prostate, bladder, gastric and endometrial cancers (41–45,40). Functionally, KNL1 is enriched in G2/M cell cycle pathways, correlates with high tumor mutation burden, microsatellite instability, T helper 2 cells (Th2)-skewed immunity, and immune checkpoint upregulation, implicating it in both chromosomal instability-driven tumor progression and immune evasion (40,46). In uterine corpus endometrial carcinoma, its silencing limits proliferation and migration of cancer cells (46). In prostate cancer, high expression correlates with regulatory T-cells (Treg) and Th2 infiltration (41).

However, despite these observations, the molecular network through which KNL1 operates and its functional relevance in HCC remain largely unexplored. Importantly, HCC is characterized by profound cell-cycle dysregulation, CIN, and immune remodelling (47), processes in which KNL1 is mechanistically positioned to play a role in. Yet, to date, up to our knowledge no study has systematically examined KNL1 within an HCC-specific, network-based and single-cell–resolved framework. Addressing this gap is critical for understanding how mitotic checkpoint dysregulation contributes to tumor progression and immune modulation in HCC.

### 1.2. Aim

This study aims to investigate the oncogenic and immunomodulatory roles of KNL1 in HCC by (1) evaluating its expression and association with chromosomal instability metrics in bulk RNA-sequence datasets from The Cancer Genome Atlas (TCGA) and Genotype-Tissue Expression (GTEx); (2) identifying and functionally annotating a KNL1-associated gene network via weighted gene co-expression network analysis (WGCNA); (3) examining the effects of this network on cell cycle distribution and TME interactions using single-cell RNA-sequence analysis; and (4) validating the biological consequences of KNL1 downregulation *in vitro* with a focus on proliferation, cell cycle regulation, and ultrastructural changes.

The strength of this work lies in its multi-modal design-combining bulk transcriptomic profiling, single-cell resolution analysis, and *in vitro* functional validation-to capture a comprehensive view of the network-level activity of KNL1. This convergence of approaches not only delineates KNL1’s molecular associations but also reveals the broader influence of its associated network on cellular behaviour, oncogenic signalling pathways, and the dynamic interplay within the TME.

## 2. Materials and Methods

The experimental and analytical procedures used in this study are summarized in ***Figure 1*** and described in detail below.

**Figure 1:**
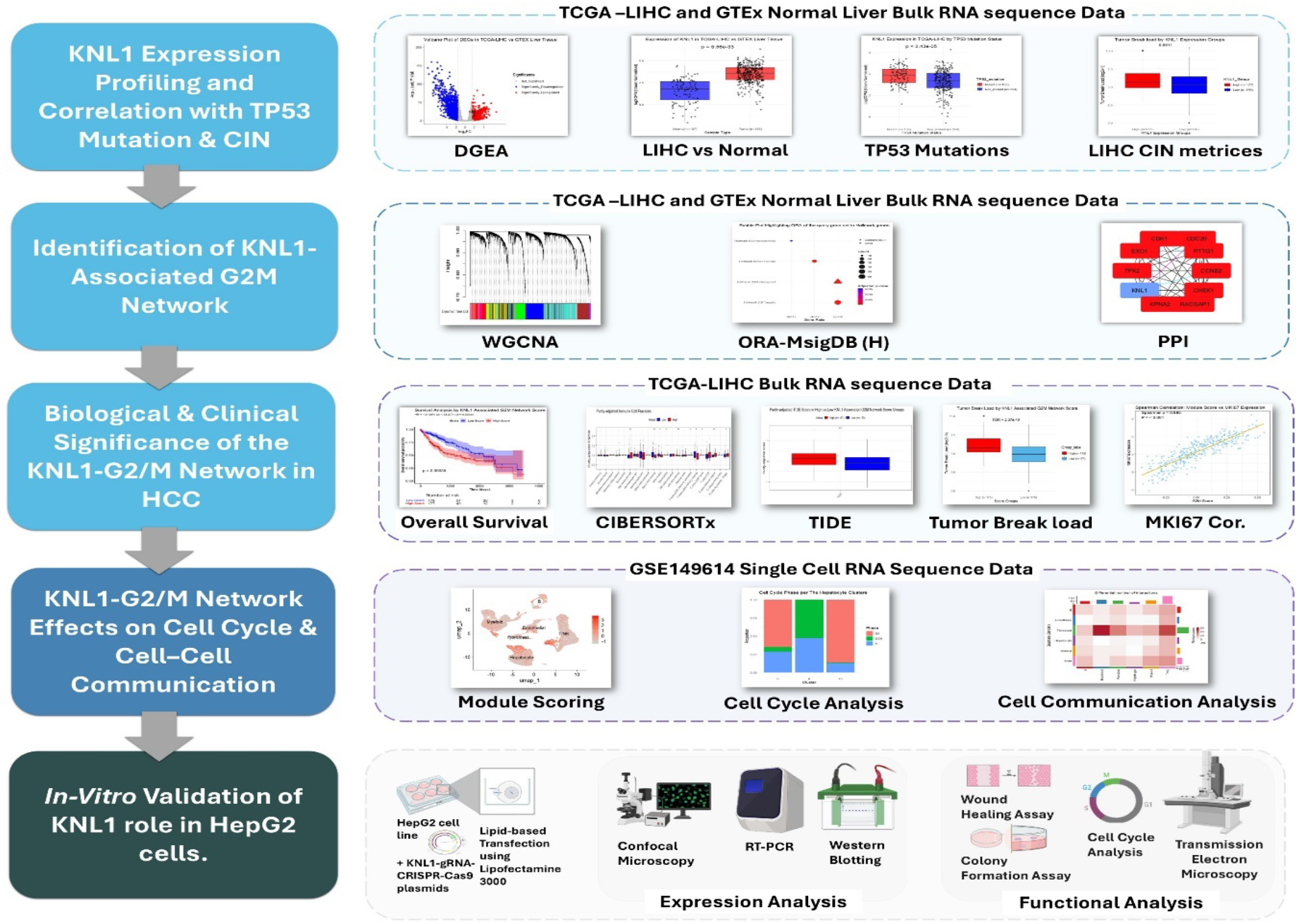
Schematic Overview of the Study Design. Step 1: Bulk RNA-seq data from TCGA-LIHC and GTEx were analysed to assess KNL1 expression, its association with TP53 mutation status, and correlation with chromosomal instability metrics, including tumor mutation burden (TMB), tumor break load (TBL), fraction genome altered (FGA) and aneuploidy score. **Step 2:** WGCNA identified a KNL1-associated gene module, which underwent functional enrichment via MSigDB hallmark gene sets and STRING protein–protein interaction network analysis**. Step 3:** Hub genes from this module were used to compute a KNL1-associated G2/M network score that was used to stratify tumors into high and low score groups for chromosomal instability (TBL, TMB, FGA and aneuploidy score), proliferation (MKI67 expression), survival, immune infiltration (CIBERSORTx) and immune evasion (TIDE scoring) analyses. **Step 4:** Single-cell RNA-seq data from GSE149614 were used to examine cell type–specific expression, cell cycle phase distribution, and cell–cell communication patterns of the KNL1-associated G2M module. Finally, functional validation was performed *in vitro* using CRISPR-Cas9-mediated downregulation of KNL1 in HepG2 cells, followed by assays for proliferation, cell cycle arrest, and ultrastructural evaluation by transmission electron microscopy (TEM). Box color codes: Light blue – Bulk RNA-seq analysis steps; Dark blue – Single-cell RNA-seq analysis; Grey – *In vitro* validation. [RNA-seq, RNA sequencing; TCGA-LIHC, The Cancer Genome Atlas Liver Hepatocellular Carcinoma; GTEx, Genotype-Tissue Expression; KNL1, Kinetochore Scaffold 1; TP53, tumor protein p53; CIN, chromosomal instability; WGCNA, Weighted Gene Co-expression Network Analysis; MSigDB, Molecular Signatures Database; STRING, Search Tool for the Retrieval of Interacting Genes/Proteins; MKI67, marker of proliferation Ki-67; CIBERSORTx, Cell-type Identification By Estimating Relative Subsets Of RNA Transcripts; TIDE, Tumor Immune Dysfunction and Exclusion; CRISPR, Clustered Regularly Interspaced Short Palindromic Repeats; Cas9, CRISPR-associated protein 9; TEM, transmission electron microscopy].

### 2.1. Bulk RNA-Sequence Analysis

#### 2.1.1. Data Acquisition and Preprocessing

Bulk RNA-seq expression data and corresponding clinical annotations (tumor stage, grade, fibrosis, and etiological risk factors) were retrieved from the UCSC Xena Toil Recompute Compendium using the UCSCXenaTools R package (48). This resource provides uniformly processed datasets from The Cancer Genome Atlas (TCGA) and Genotype-Tissue Expression (GTEx) via the Toil pipeline to minimize technical batch effects (49). We analysed primary HCC tumors from TCGA-liver hepatocellular carcinoma cohort (LIHC) (n=359) against histologically normal liver tissues from GTEx (n=110) using RNA-Seq by Expectation-Maximization counts (RSEM).

Additional clinical features-including aneuploidy score, tumor break load, tumor mutation burden and fraction genome altered-were retrieved from cBioPortal (https://www.cbioportal.org) (LIHC TCGA PanCan Atlas 2018 dataset) (50).

Tumor protein p53 (TP53) mutation status was obtained from the Multi-Center Mutation Calling in Multiple Cancers (MC3) dataset (<u>mc3.v0.2.8.PUBLIC.toil.xena.gz</u>) hosted on the UCSC Xena platform (https://xenabrowser.net/datapages/). The gene annotation file (zz_gencode.v23.annotation.csv) used for ID–gene symbol mapping was downloaded from the UCSC Xena website (<u>gencode.v23.annotation.gene.probemap</u>), and only protein-coding genes were retained for downstream analyses. Preprocessing steps using Bioconductor workflows were partially adapted from a published protocol (51), with modifications optimized for our study.

#### 2.1.2. Differential Gene Expression Analysis

To identify genes with significant transcriptional alterations between HCC and normal liver tissue, differential gene expression analysis (DGEA) was performed using an edgeR–limma–voom pipeline (52–54). Quality control was performed using density plots and principal component analysis (PCA). Subsequently, a DGEList object was constructed in edgeR, and genes with low expression across all samples were filtered. To correct for library size and composition bias, counts were normalized using the upper quartile (UQ) method. Voom transformation was then applied to model the mean–variance relationship and assign observation-level precision weights.

A linear design matrix contrasting tumor (n= 358) versus normal tissue (n= 107) was used for model fitting, followed by empirical Bayes moderation. Differentially expressed genes (DEGs) were defined using the following thresholds (absolute Log₂ fold-change (|log_2_FC|) ≥ 1.5, Benjamini–Hochberg adjusted p-value < 0.05). Results were visualized using volcano plot highlighting KNL1 gene.

#### 2.1.3. KNL1 Expression Analysis

To investigate the relationship between KNL1 expression and chromosomal instability, TCGA-LIHC tumor samples were stratified into high and low KNL1 expression groups based on the median KNL1 expression value. Chromosomal instability-related metrics—including tumor break load, tumor mutation burden, aneuploidy score and fraction genome altered as obtained from cBioPortal—were compared between groups using the Wilcoxon rank-sum test (50,55,56).

Additionally, somatic mutation data from the MC3 dataset were filtered to include only TCGA liver tumor samples. Samples harbouring disruptive TP53 mutations (e.g., missense, nonsense, frameshift) were classified as mutant, other samples including those with only silent, intronic, and wild type were categorized as non-mutant (57). KNL1 expression was compared between these groups using the Wilcoxon rank-sum test.

#### 2.1.4. Weighted Gene Co-expression Network Analysis (WGCNA)

Weighted gene co-expression network analysis (WGCNA) using WGCNA R package (version 1.73) was performed to identify gene modules that share expression pattern similarity with KNL1 in HCC (58,59). Voom-transformed expression data from TCGA-LIHC tumor samples were filtered to retain the top 75% most variable genes, and outlier samples were excluded based on hierarchical clustering of sample distances.

A soft-thresholding power of 10 was selected to approximate scale-free topology (R² ≈ 0.90), and a signed topological overlap matrix (TOM) was computed. Gene clustering was performed using average linkage hierarchical clustering of TOM-based dissimilarity, and modules were detected using dynamic tree cutting (hybrid method) with minimum module size of 70 genes and Deep split 4, allowing for high sensitivity in detecting distinct gene modules. Module eigengenes (MEs) were calculated and correlated with clinical traits to identify biologically meaningful modules using Pearson correlation. Within each module, module membership (MM) scores were computed as the correlation between gene expression and module eigengene values. Final gene-module assignments and annotations were exported for downstream functional enrichment and network analysis.

#### 2.1.5. Over-Representation Analysis (ORA)

To refine the list of KNL1-associated genes, significantly upregulated genes were identified using the criteria |log_2_FC| > 1.5 and Benjamini–Hochberg adjusted p-value < 0.05. Genes overlapping between this gene set and the KNL1-associated green module identified via WGCNA—were selected for downstream pathway over-representation analysis (ORA).

Gene symbols were first converted to Entrez Gene IDs using the org.Hs.eg.db package to enable compatibility with enrichment tools (60). ORA was conducted using the enricher() function from the clusterProfiler R package (version 4.16.0) (61–63). The Hallmark (H) gene set was obtained from the Molecular Signatures Database (MSigDB) via the msigdbr package (version 25.1.1) (64,65). Enriched terms with Benjamini–Hochberg adjusted p-value < 0.05 were considered statistically significant.

The results were visualized using bubble plot to display gene ratios, and adjusted p-values. Among the significantly enriched terms, particular attention was given to the G2M checkpoint hallmark, in which KNL1 was found to be included.

#### 2.1.6. Protein–Protein Interaction (PPI) and G2M Hallmark Gene Analysis

PPI networks were constructed using Cytoscape (v3.10.3) and the stringApp plugin, based on gene sets derived from the intersection of the KNL1-associated WGCNA module, significantly upregulated DEGs, and the Hallmark G2M checkpoint gene set obtained from MSigDB (66,67). This intersection was visualized using a venn diagram generated with the ggvenn R package (68). The Search Tool for the Retrieval of Interacting Genes/Proteins (STRING) database was queried directly within Cytoscape, using a minimum confidence score of 0.15 to include both high- and moderate-confidence interactions. The resulting network visualized proteins as nodes and their functional associations as edges (67,69).

log₂FC values from the differential expression analysis were imported into the Cytoscape node attribute table and mapped to node color, with all nodes rendered in shades of red since the intersected gene set consisted exclusively of significantly upregulated genes. The intensity of red was scaled to represent the magnitude of log₂FC, allowing a visual distinction between moderately and highly upregulated genes. Network topology was analysed using Cytoscape’s built-in Network Analyzer tool, and a circular layout was applied for visualization. The annotated node table—including topological metrics was then exported for downstream analysis. Hub genes were identified using the cytoHubba plugin. The Maximal Clique Centrality (MCC) algorithm was applied to the STRING-derived network, and the top 10 genes ranked by MCC score were extracted for subsequent biological interpretation and validation (70).

#### 2.1.7. Single Sample Gene Set Enrichment Analysis (ssGSEA) and Score Stratification

To avoid incorporating genes with lower connectivity or weaker relevance to KNL1-centered regulation, which could dilute signals specific to the KNL1-associated network, the top 10 hub genes identified by MCC scoring were selected for ssGSEA-based pathway activity estimation. ssGSEA was performed using the GSVA R package (version 3.21) (method = “ssgsea”) on voom-transformed expression data from TCGA-LIHC tumor samples, yielding a continuous enrichment score for each sample, referred to as the “KNL1-Associated G2M Network Score” (71).

Samples were then stratified into “High” and “Low” score groups based on the median enrichment score to facilitate downstream comparisons. Alternative stratification approaches, including quartile-based grouping, were tested during method development and yielded consistent results in immune and chromosomal instability analyses; therefore, the median cutoff was retained for consistency across analyses.

In parallel, expression data for the whole 60-gene Hallmark G2M checkpoint signature—previously identified via over-representation analysis (ORA)—were extracted, Z-score normalized, and visualized using the ComplexHeatmap package (72). Heatmaps were annotated based on both KNL1-associated network score groups (high vs. low) and sample type (tumor vs. normal) to aid in the interpretation of expression patterns across biological contexts.

#### 2.1.8. Survival, Proliferative Potential, and Chromosomal Instability Analysis

To evaluate the prognostic relevance of KNL1 and its associated proliferative network, KNL1-related gene set scores were integrated with TCGA clinical annotations. Samples with overall survival (OS) exceeding 10 years (>3,650 days) were excluded. Survival analyses were conducted using Kaplan–Meier (KM) estimation and Cox proportional hazards regression implemented in the *survival* and *survminer* R packages, with statistical significance assessed using the log-rank test (73,74).

For survival visualization, patients were dichotomized into high- and low-risk groups based on the median single-sample gene set enrichment analysis (ssGSEA) score of the KNL1-associated G2M network, and KM survival curves were generated accordingly. This dichotomized grouping was applied for graphical representation and group-wise comparison. For risk estimation and inference, the G2M network score was scaled and modelled as a continuous variable. Univariate and multivariate Cox proportional hazards regression analyses were performed to assess its association with OS and to determine whether the score represents an independent prognostic factor. Multivariate models were adjusted for clinically relevant covariates, including age at diagnosis, sex, histological grade, pathological stage, vascular invasion, serum alpha-fetoprotein and viral hepatitis status. The predictive performance of the KNL1-associated G2M ssGSEA score for overall survival (OS) at 1-, 3-, and 5-year time points using time-dependent ROC curves with inverse probability of censoring weighting (IPCW) (75).

To further contextualize pathway-level findings, we also evaluated the association between KNL1 expression alone and OS using both median-based and quartile-based stratification approaches.

#### 2.1.9. External Validation of Prognostic Relevance of KNL1-associated G2M Network

The prognostic robustness of the KNL1-associated G2M gene network was further evaluated in independent external cohorts, including the Liver Cancer—RIKEN, Japan Project from the International Cancer Genome Consortium ((http://platform.icgc-argo.org; ICGC LIRI-JP) (76) and the microarray GEO datasets GSE54236 and GSE14520 (77,78). For the LIRI-JP cohort, gene expression values quantified as fragments per kilobase million (FPKM) were first converted to transcripts per million (TPM) and log₂-transformed (log₂[TPM + 1]) prior to downstream analyses. For GSE54236 and GSE14520, normalized expression matrices were obtained directly from the Gene Expression Omnibus (GEO) using GEOquery R package (79). The 10-gene KNL1-associated G2M ssGSEA score was calculated for each sample using the *GSVA* package, followed by scaling and integration with available clinical metadata (71). In GSE14520, analyses were restricted to tumor samples profiled on the GPL3921 platform to ensure platform consistency.

In LIRI-JP and GSE1450, for survival visualization, tumor samples were dichotomized into high- and low-risk groups based on the median G2M ssGSEA score, and Kaplan–Meier survival curves were generated with statistical significance assessed using the log-rank test. This dichotomized grouping was applied exclusively for visualization. For prognostic inference, the G2M network score was modeled as a continuous variable in Cox proportional hazards regression analyses. Univariate Cox models were first fitted to assess the association between the G2M score and OS. Multivariate Cox models were then constructed to evaluate the independence of the G2M score after adjustment for cohort-specific clinical covariates. In GSE14520, multivariate models were adjusted for age, sex, serum alpha-fetoprotein (AFP), and hepatitis B virus (HBV) infection status, whereas in LIRI-JP, adjustments included age, sex, tumor grade, and clinical stage. Predictive performance and discriminatory ability were further assessed using time-dependent receiver operating characteristic (ROC) analyses at 1-, 3-, and 5-year endpoints (75).

In GSE54236, complete survival time and censoring information were not available. Therefore, a fixed 1-year OS endpoint was defined (survival ≥12 months = 0 (alive); survival <12 months = 1 (deceased)). Prognostic associations in this cohort were evaluated using Cox regression applied to the fixed 1-year endpoint, as well as logistic regression modeling 1-year survival as a binary outcome using the scaled G2M ssGSEA score as a continuous variable to preserve quantitative information.

#### 2.1.10. Proliferative capacity and Chromosomal Instability Analysis

To assess proliferative potential, Spearman’s correlation between KNL1-associated network scores and marker of proliferation Ki-67 (*MKI67)* expression in tumor samples was computed and visualized using ggplot2 (80). Chromosomal instability–related metrics, including tumor break load, tumor mutation burden, aneuploidy score, and fraction genome altered, were retrieved from cBioPortal and compared between high- and low-score groups using the Wilcoxon rank-sum test and p values were corrected for multiple testing using Benjamini–Hochberg method (50,55,56).

#### 2.1.11. Immune Infiltration and Immune Evasion Analysis

Tumor immune cell composition was estimated using CIBERSORTx, which deconvolutes bulk RNA-seq data to infer the relative abundance of 22 immune cell types. TCGA liver cancer (LIHC) data were obtained from the PanImmune dataset (TCGA.Kallisto.fullIDs.cibersort.relative.tsv; https://gdc.cancer.gov/about-data/publications/panimmune) (81,82). Tumor purity estimates were retrieved from the TCGA PanCancer Atlas (TCGA_mastercalls.abs_tables_JSedit.fixed.txt; https://gdc.cancer.gov/about-data/publications/pancanatlas) and used to adjust downstream analyses. Functional immune evasion was assessed with the TIDE framework, quantifying overall TIDE score, T-cell dysfunction and T-cell exclusion (83). Log₂(TPM + 0.001)-transformed TCGA-LIHC expression data were pre-processed for TIDE input.

CIBERSORT immune fractions, tumor purity, and TIDE scores were integrated with KNL1-associated network scores. Samples were stratified into high- and low-score groups using the median network score. To account for tumor purity, TIDE scores (Overall TIDE, Exclusion, Dysfunction) and immune fractions were first residualized via linear regression (84–86):

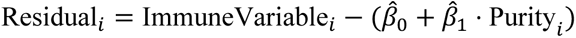

where *β̂*_0_ and *β̂*_1_ are the intercept and slope from regressing the immune variable on tumor purity. Immune comparisons between high- and low-score groups were performed using Wilcoxon rank-sum tests, with Benjamini–Hochberg correction for multiple testing.

### 2.2. Single-Cell RNA-Seq Analysis

#### 2.2.1. Data Processing

GSE149614 scRNA-seq data (71,915 cells) from HCC and adjacent tissues were processed using Seurat (version 5.3.0), quality-controlled, and integrated across samples using canonical correlation analysis (CCA) to assess and correct batch effects (87–90). Data were analysed via PCA, Uniform Manifold Approximation and Projection (UMAP), and Louvain clustering (resolution = 0.3) to capture major cell type groupings. Cells were annotated based on the original metadata, and cell type distributions were visualized as stacked bar plots depicting both proportional and absolute counts (91).

#### 2.2.2. Module Scoring and Cell Cycle Analysis

To evaluate KNL1-associated G2M network activity at single-cell resolution, we applied AddModuleScore() Seurat function using the full 60-gene Hallmark G2M checkpoint signature, including KNL1. Although 10 hub genes were prioritized in bulk analysis based on network centrality, the full gene set was used in single cell analysis to enhance biological coverage and mitigate dropout effects inherent to single-cell RNA-seq (92).

Cell cycle phases (G1, S, G2/M) were assigned using Seurat’s CellCycleScoring() function. Hepatocyte-enriched clusters (0, 8, 15) were selected to assess the relationship between KNL1-associated G2M module scores and cell cycle state.

#### 2.2.3. Pseudo-bulk Analysis

Single-cell UMI counts were aggregated per patient × cluster to generate pseudobulk profiles. Samples with <100 cells or <1×10⁴ UMIs were removed. Counts were imported into edgeR, lowly expressed genes were filtered (CPM > 1 in ≥2 samples), and normalization was performed using TMM.

Pathway activity was quantified using ssGSEA (GSVA) for three gene sets: the KNL1-associated G2M module, Seurat S-phase genes, and Seurat G2/M-phase genes (obtained from Seurat; cc.genes.updated.2019). Only patients contributing both clusters were used for paired testing. Differences in ssGSEA scores between clusters (e.g., cluster 8 vs 0) were assessed using paired Wilcoxon tests.

Differential gene expression between pseudobulk clusters was performed using limma-voom with patient blocking (∼ patient + cluster).

#### 2.2.4. Cell-Cell Communication Analysis

Tumor cells were stratified based on both the KNL1-associated G2M module score (median score/cell type) and KNL1 expression (positive vs. negative). Cells with high KNL1-associated G2M module score and positive KNL1 expression were grouped as “High_Positive” while those with low KNL1-associated G2M module score and negative KNL1 expression were grouped as “Low_Negative”. These dual-stratified groups were used for subsequent CellChat analysis (93,94).

CellChat objects were constructed separately for each group and processed via the standard pipeline: identification of overexpressed genes and ligand–receptor interactions, computation of communication probability (min.cells = 10), pathway inference, and network aggregation. Comparative analyses included netVisual_heatmap() and rankNet() to assess global signalling changes between groups.

To visualize directional intercellular communication patterns, we applied netVisual_bubble() to merged CellChat objects comparing High_Positive and Low_Negative tumor groups. ECM-related signalling pathways—including LAMININ, ANGPTL, TENASCIN, and FN1—were selected based on their significant enrichment in High_Positive tumors and visualized to examine ligand–receptor interactions between hepatocytes and fibroblasts. Similarly, immune-modulatory pathways such as MIF, CXCL and MHC-I were selected for visualization based on differential enrichment, enabling assessment of bidirectional immune crosstalk between tumor/stromal and immune cell populations (B cells, T/NK cells, and myeloid cells).

### 2.3. *In Vitro* Validation of KNL1 Function

#### 2.3.1. Materials and reagents

Human liver hepatocellular carcinoma cell line (HepG2) cell line used in this work was obtained from the American Type Culture Collection (ATCC) (https://www.atcc.org/). Antibodies against KNL1 (#PA5-88989) were from (Thermofisherscientific, USA). Dulbecco’s Modified Eagle Medium (DMEM), fetal bovine serum (FBS), penicillin/streptomycin and Trypsin/EDTA (Invitrogen, USA). KNL1 CRISPR guide RNA_pSpCas9 BB-2A-GFP-PX458 vectors were purchased from (Genscript, USA) to obtain all in one plasmid containing SpCas9 enzyme, KNL1 gRNA and GFP reporter. Guide RNA (gRNA) sequences were ATCGAACCAAGATAAGCTGT and AATGGATGGGGTGTCTTCAG for gRNA1 and gRNA2 respectively.

#### 2.3.2. Cell Culture

HepG2 cells were cultured in DMEM medium containing 10% fetal bovine serum (FBS), 1% penicillin-streptomycin at 37 °C with 5% CO_2_. The cells were subcultured with trypsin-EDTA when they reached 70–80% confluency.

#### 2.3.3. Transfection of HepG2 Cells

CRISPR-Cas9–mediated dual targeting of KNL1 was employed in HepG2 cells, as previous studies indicate that using multiple gRNAs at lower concentrations can improve editing efficiency (95–97). HepG2 cells were seeded in 6-well plates at a density of 5 × 10⁵ cells/well and cultured for 24 hours prior to transfection. Cells in the KNL1 gRNA1+2 group were co-transfected with plasmids carrying KNL1-targeting gRNA1 and gRNA2 using Lipofectamine 3000 (Invitrogen, USA) in Opti-MEM I Reduced Serum Medium (Gibco, USA). After 72 hours, cells were harvested for downstream analyses. Transfection efficiency was evaluated by GFP reporter expression, while KNL1 knockdown efficiency was confirmed by real-time quantitative PCR (RT-qPCR) and Western blotting.

#### 2.3.4. Confocal Microscopy

The KNL1 targeting plasmids were transfected into HepG2 cells in 35 mm cell culture dish (NEST, USA) according to the manufacturer’s instructions. After 72 h of transfection, cells were examined under a confocal microscope for GFP fluorescence using Leica DMi8 A (Wetzlar, Germany) and analysed using Leica Application Suite X (LAS X, core software v 3.7.4; Leica). The mean fluorescence intensity (MFI) was analysed by imageJ in randomly captured fields to quantify GFP expression (98).

#### 2.3.5. RNA Isolation and Real-Time Quantitative PCR (RT-qPCR)

Real-time quantitative PCR (RT-qPCR) was performed to quantify the mRNA expression of KNL1 in both KNL1 gRNA1+2 group and the control group. To further investigate the impact of KNL1 downregulation on the pro-apoptotic response, the expression levels of BIRC5 (survivin) and BCL2 were also assessed (99,100).

Total RNA was extracted using TRIzol reagent (Invitrogen, USA), and reverse transcribed into cDNA using the RevertAid First Strand cDNA Synthesis Kit (Thermo Scientific, USA), following the manufacturer’s instructions.

RT-qPCR was carried out using Maxima SYBR Green/ROX qPCR Master Mix (2X) (Thermo Scientific, USA) on a StepOnePlus Real-Time PCR System (Applied Biosystems, USA). The reaction employed a two-step thermal cycling protocol: initial denaturation at 95 °C for 10 min, followed by 45 cycles of 95 °C for 15 s and 60 °C for 1 min. Gene expression levels were normalized to GAPDH as an internal control, and fold changes were calculated using the 2^-ΔΔCt method. The primer sequences used for KNL1 were: forward 5′-ACCCTGAAGTCACCATCTGT-3′ and reverse 5′-GGTTGGGGAACATCACTCAC-3′.

#### 2.3.6. Western Blot Analysis

HepG2 cell pellets were resuspended in homogenization buffer supplemented with protease & phosphatase inhibitor cocktail (Thermofisherscientific, USA) and sonicated on ice (30% amplitude-for 2 × 5 sec). Sonicates were centrifuged at 4°C (10,000×g for 5 min). Supernatants were collected for protein determination using nanodrop spectrophotometer (Thermo, USA). Lysate proteins (50 µg) were loaded onto 4-12% polyacrylamide gel and separated by electrophoresis using a Biorad Tetra Cell (Biorad, USA) for 90 min at 150 volts. Resolved proteins were transferred to nitrocellulose membrane by blotting using a Biorad Trans blot module for 75 minutes at 100 volts under cooling. Membranes were blocked using 5% non-fat milk in Tris buffered saline tween 20 (TBST) for 1.5h at room temperature (∼25 °C), with gentle shaking and probing with primary KNL1 antibody (1:1000) or β-actin (1:2000) (Thermofisherscientific, USA). Membranes were incubated with antibodies at 4°C overnight with gentle rolling on a tube roller. On the next day, membranes were washed 3 × 5 min with TBST. Membranes were then probed with the corresponding horseradish peroxidase (HRP)-conjugated secondary antibodies for 1 h at room temperature (∼25 °C). The membranes were washed as above with TBST. Enzyme chemiluminescence (Pierce, Thermofisherscientific, USA) was used to visualize the protein bands. The bands were captured using a UVP Biospectrum Imager. Band intensities were quantified using ImageJ software and normalized to those of β-actin (98).

#### 2.3.7. Wound-Healing Assay

The cell migration potential following transfection was assessed by the monolayer wound healing assay. After 72 hr of transfection, and a straight line across wells of six-well plate was drawn using a sterile 100 μL pipette tip. The scratched cells were washed off with phosphate-buffered saline (PBS) and captured at 0, 24 and 48 h using a ZEISS Primovert inverted microscope. Wound areas were measured using ImageJ software (National Institutes of Health, Bethesda, MD, USA; https://imagej.net/ij/) and the percentage of wound closure at each time point was calculated relative to the initial wound area at 0 hours (98).

#### 2.3.8. Colony Formation Assay

HepG2 cells were washed with PBS and trypsinised. Cells were then counted and a total of 2,000 HepG2 cells from each group (Control and Treatment) were seeded into 6-well plates and incubated at 37 °C with 5% CO₂ in a humidified incubator. After 10 days of culture, the medium was discarded, and wells were gently rinsed with PBS. Cells were fixed with a 1:7 acetic acid: methanol solution for 5 minutes at room temperature (∼25 °C), followed by staining with 0.2% crystal violet for 30 minutes. After thorough washing, colonies were captured and quantified using the colonyArea plugin of ImageJ software (101).

#### 2.3.9. Cell Cycle Analysis

Cells (1 × 10⁶) were suspended in 0.5 ml of 1× Dulbecco’s phosphate-buffered saline (DPBS) and gently aspirated several times using a Pasteur pipette. The cells were then fixed in 70% ethanol on ice for 2 hours. Following fixation, cells were centrifuged at 300 ×g for 5 minutes and the ethanol was thoroughly removed. The cell pellets were washed by resuspension in 5 ml of 1× DPBS for 30 seconds, followed by centrifugation at 300 ×g for 5 minutes. Pellets were then resuspended in 1 ml of propidium iodide (PI) staining solution and incubated in the dark at room temperature (∼25 °C) for 30 minutes. Fluorescence was measured using a CytoFLEX Flow Cytometer (Beckman Coulter Life Sciences, USA). The distribution of cells across the G1, S, and G2/M phases was analysed using CytExpert Software (Beckman Coulter Life Sciences, USA; https://www.beckman.com/flow-cytometry/software/cytexpert).

#### 2.3.10. Transmission Electron Microscopy (TEM)

Cells were fixed in 2.5% glutaraldehyde for 4 hours at 4 °C, followed by three washes in PBS. Post-fixation was performed using 1% osmium tetroxide (osmic acid) for 1 hour at room temperature (∼25 °C). Samples were then washed and dehydrated through a graded ethanol series. Dehydrated specimens were cleared in acetone and embedded in epoxy resin. Ultrathin sections (60–90 nm) were cut using an ultramicrotome (RMC, Boeckeler Instruments; USA) and mounted on copper grids. Sections were subsequently stained with uranyl acetate and lead citrate. Finally, they were examined using transmission electron microscopy (JEM 1200EX II, Jeol; Japan) operated at an accelerating voltage of 80 kV.

### 2.4. Statistical analysis

All analyses were performed in R (version 4.5.1). Data distribution was assessed using normality testing (Shapiro–Wilk), and statistical tests were selected accordingly. RNA-seq–derived measures, gene set scores, chromosomal instability metrics, and immune cell fractions did not meet normality assumptions and were therefore treated as non-parametric; group comparisons were performed using Wilcoxon rank-sum or signed-rank tests, and results are reported as median ± interquartile range (IQR). Experimental data from in vitro assays were normally distributed and analyzed as parametric, using two-tailed Student’s *t*-tests, and are presented as mean ± standard deviation (SD). Correlations were assessed using Spearman’s rank correlation. Survival analyses were conducted using Kaplan–Meier methods with log-rank tests and Cox proportional hazards models, adjusting for relevant clinical covariates. Multiple testing correction was applied using the Benjamini–Hochberg false discovery rate (FDR) for bulk and pseudobulk RNA-seq analyses. Statistical significance was defined as *p* < 0.05.

## 3. Results

### 3.1. KNL1 is Overexpressed in HCC and Associated with Genomic Instability

After running PCA, 4 samples (“TCGA.UB.AA0V.01”, “GTEX.O5YT.0826.SM.3TW8N”, “GTEX.WYVS.1926.SM.4PQZ2” and “GTEX.1192X.1026.SM.5H12P”) were detected as outliers and removed before downstream analysis. PCA showed clustering of samples based on the condition (tumor versus normal) (***Figure 2A***). Differential gene expression analysis revealed 636 significantly upregulated and 1695 significantly downregulated genes (|log_2_FC| > 1.5 and *adj.p* < 0.05). Among the significantly upregulated genes, KNL1 was markedly overexpressed in HCC primary tumor tissues compared to normal liver control samples (Log₂FC = 1.9, *adj. p* = 4.33E-25) (***Figure 2B***).

**Figure 2:**
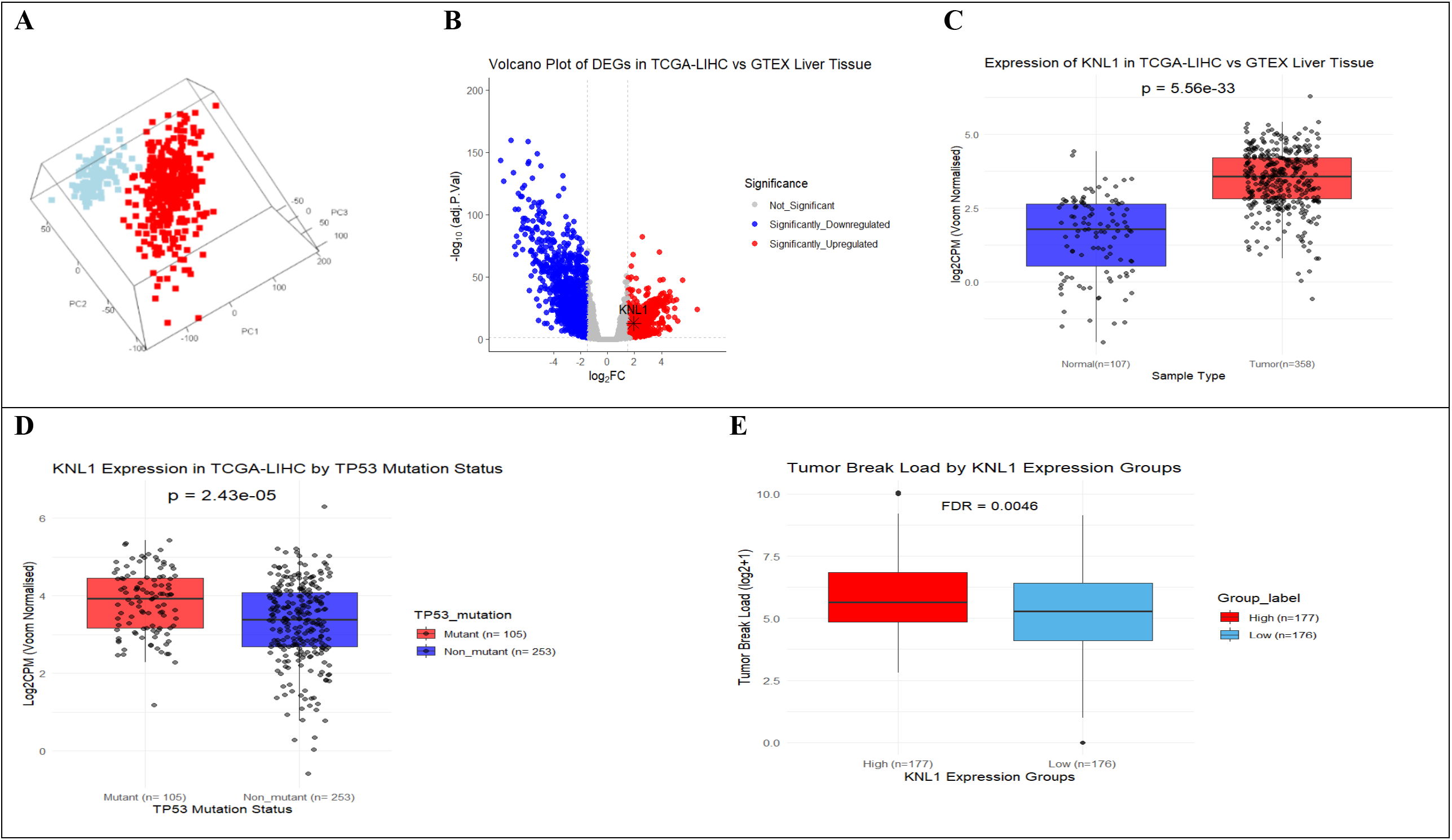
KNL1 is upregulated in HCC and linked to chromosomal instability. (**A**) Principal component analysis (PCA) demonstrated a clear separation between tumor (n = 358) and normal liver samples (n = 107) following the exclusion of four outliers. (B) Volcano plot illustrating differentially expressed genes, identifying 636 upregulated and 1,695 downregulated genes using thresholds of |log₂ fold change| > 1.5 and adjusted *p* < 0.05. KNL1 was significantly overexpressed in tumor samples (log₂FC = 1.9, adjusted *p* = 4.33 × 10⁻²⁵). **(C)** Boxplot analysis confirmed significantly higher KNL1 expression in TCGA-LIHC tumors compared to normal GTEx liver tissues (median [IQR]: 3.56 [2.81–4.20] vs. 1.79 [0.53–2.63], median Δ= 1.77, adjusted *p* = 5.56 × 10⁻³³). **(D)** KNL1 expression was further elevated in TP53-mutant tumors (n= 105) compared with TP53 non-mutant group (n=253) (median [IQR]: 3.92 [3.16–4.45] vs. 3.38 [2.69–4.08], median Δ= 0.54, adjusted *p* = 2.43 × 10⁻⁵). (E) High KNL1 expression was associated with increased tumor break load (median [IQR]: 49 [28–115] vs. 37.5 [16–84.5], median Δ= 11.5, adjusted *p* = 0.0046), indicating a link between KNL1 and structural chromosomal instability. The y-axis represents log-transformed TBL values to facilitate visualization of the data distribution. For panels (C–D), statistical comparisons were performed using the Wilcoxon rank-sum test. Data are presented as median and interquartile range (IQR). Adjusted *p* values were calculated using the Benjamini–Hochberg method, with adjusted *p* < 0.05 considered statistically significant. [TCGA-LIHC, The Cancer Genome Atlas Liver Hepatocellular Carcinoma; GTEx, Geno-Type Tissue Expression; PCA, Principal component analysis].

Boxplot analysis confirmed a significant upregulation of KNL1 expression in TCGA-LIHC primary tumor samples compared with normal GTEx liver tissues (median [IQR]: 3.56 [2.81–4.20] vs. 1.79 [0.53–2.63], median Δ= 1.77, *adj. p* = 5.56 × 10⁻³³; ***Figure 2C***). Similarly, **KNL1** expression was significantly higher in TP53-mutant primary tumors than in TP53 wild-type samples (median [IQR]: 3.92 [3.16–4.45] vs. 3.38 [2.69–4.08], median Δ= 0.54, *adj. p* = 2.43 × 10⁻⁵; ***Figure 2D***).

After stratifying tumors into high- and low-KNL1 expression groups based on the median KNL1 expression, boxplot analysis revealed that tumors in the high-KNL1 group exhibited a significantly higher tumor break load (TBL) compared with those in the low-KNL1 group (median [IQR]: 49 [28–115] vs. 37.5 [16–84.5], median Δ= 11.5, *adj. p* = 0.0046), suggesting a potential association between KNL1 expression and chromosomal instability in HCC. For improved visualization of the wide distribution of values, TBL was log-transformed in the corresponding ***Figure 2E***. In contrast, no significant differences were observed between high- and low-KNL1 expression groups with respect to aneuploidy score (median [IQR]: 8 [5–14] vs. 8 [5–19], median Δ= 0, adjusted *p* = 0.619), fraction of genome altered (0.260 [0.178–0.381] vs. 0.239 [0.148–0.378], median Δ= 0.021, *adj. p* = 0.20), or tumor mutation burden (2.90 [2.19–3.98] vs. 2.88 [1.92–4.07], median Δ= 0.02, *adj. p* = 0.619) ***(Supplementary Figures S1A–C)*.**

### 3.2. KNL1 is a Central Regulator of the G2M Checkpoint Network in HCC

#### 3.2.1. Weighted Gene Co-Expression Network Analysis (WGCNA)

Using WGCNA, 15 gene co-expression modules were identified through hierarchical clustering and dynamic tree cutting (***Figure 3A***). Of these, the green module showed the most prominent and statistically significant association with KNL1 expression (Module Membership = 0.427, *p* = 1.37E-16).

**Figure 3:**
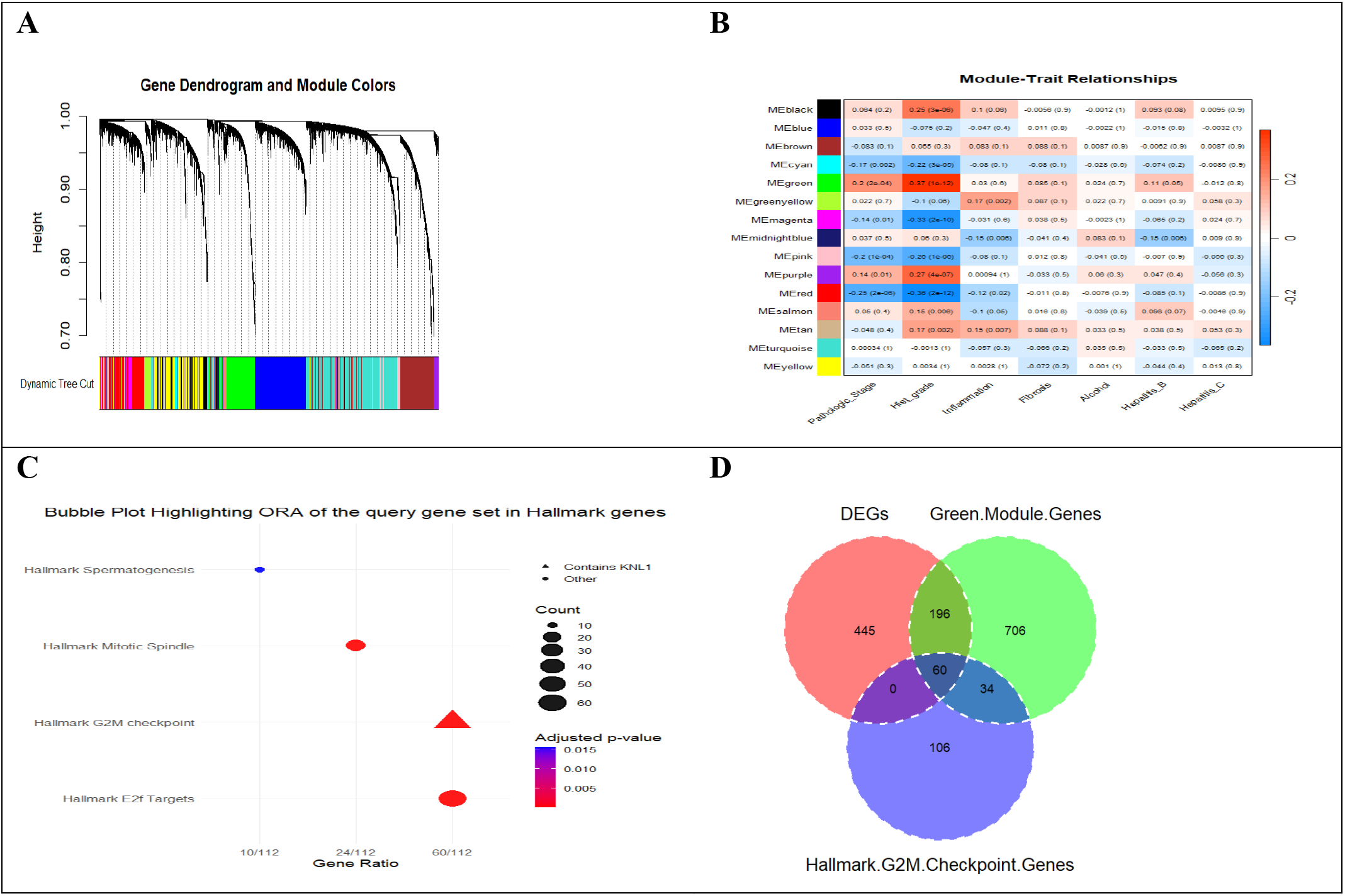
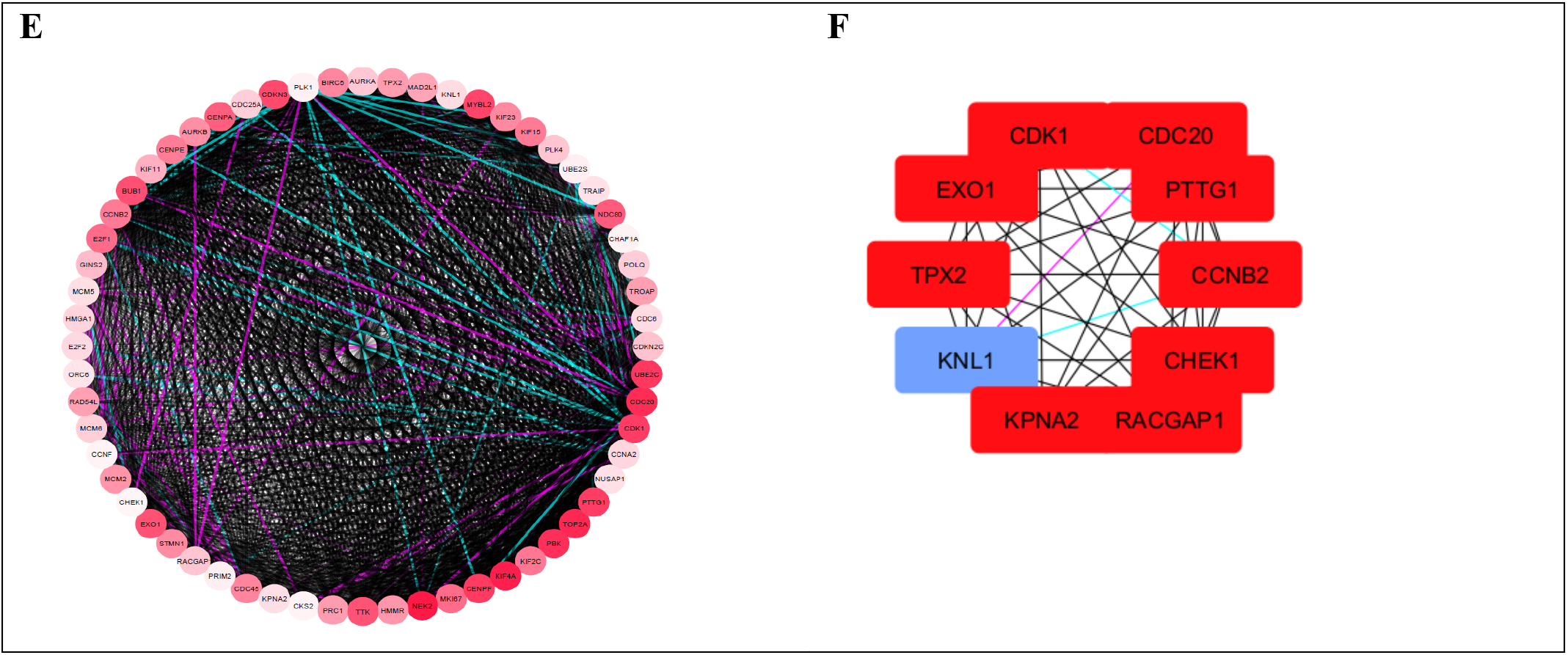
WGCNA-based identification and functional characterization of the KNL1-associated module in HCC. (**A**) Gene dendrogram showing 15 co-expression modules identified by WGCNA using hierarchical clustering and dynamic tree cutting. (**B**) Module–trait heatmap showing that the green module has the strongest positive correlation with histological grade (ρ = 0.37, *p* = 1E-12). (**C**) Bubble plot of Hallmark pathway enrichment for the 256 differentially expressed genes (DEGs) in the green module, highlighting enrichment in G2M checkpoint, E2F targets, and mitotic spindle pathways. Bubble size indicates gene count, and color reflects the adjusted p-value. (**D**) Venn diagram showing 60 overlapping genes among the green module, significantly upregulated DEGs, and the Hallmark G2M Checkpoint gene set. (**E**) Protein–protein interaction (PPI) network of the 60 intersecting genes constructed using STRING in Cytoscape. The network consists of 60 nodes and 1,708 edges, with a high average node degree (56.93), clustering coefficient (0.975), and density (0.965), indicating a highly interconnected and co-regulated structure. Node color indicates log₂ fold change (log₂FC), with deeper red shades representing higher levels of upregulation. Edge colors represent interaction evidence types: cyan (databases), light green (text mining), pink (experiments), black (co-expression), green (neighbourhood), blue (co-occurrence), and red (gene fusion). (**F**) Top 10 hub genes identified using Maximal Clique Centrality via cytoHubba, including KNL1, among central regulators of G2/M checkpoint signalling. [WGCNA, weighted gene co-expression network analysis; STRING, Search Tool for the Retrieval of Interacting Genes/Proteins; cdc20, cell division cycle protein 20 homolog; PTTG1, Pituitary tumor transforming gene 1; CCNB2, Cyclin B2; CHEK1, checkpoint kinase 1; RACGAP1, Rac GTPase Activating Protein 1; KPNA2, karyopherin subunit alpha 2; KNL1, Kinetochore scaffold 1; TPX2, Targeting protein for Xenopus kinesin-like protein 2; EXO1, Exonuclease 1; CDK1, cyclin dependent kinase 1].

Moreover, the green module showed the strongest positive correlation with histological grade (ρ = 0.37, *p* = 1E-12), as visualized in the module–trait heatmap (***Figure 3B***). This suggests that the genes within this module may play a role in tumor differentiation and progression. Further analysis revealed a strong positive correlation between module membership (MM) and gene significance (GS) for histological grade (cor = 0.75, *p* = 1.2e-180). Additionally, moderate correlations were observed between MM and GS for pathological stage (cor = 0.62, *p* =7.8 e-107) and fibrosis (cor = 0.42, *p* =7e-44), highlighting the potential involvement of green module genes in HCC tumorigenesis ***(Supplementary Figure S2)*.**

#### 3.2.2. Functional Enrichment of the KNL1 Module Genes

Intersection of the genes from the KNL1-associated green module obtained by WGCNA with the differentially expressed genes (DEGs), generated a refined list of 256 DEGs within the green module. This intersected gene set served as the input for over-representation analysis (ORA).

Using the MSigDB Hallmark gene sets, enrichment analysis revealed that our query gene set was significantly enriched in E2F Targets, G2M Checkpoint, Mitotic Spindle, and Spermatogenesis pathways.

***Table 1*.** summarizes each enriched hallmark pathway along with the corresponding gene count and gene identifiers. Notably, KNL1 was among the 60 genes enriched in the G2M Checkpoint hallmark, with a fold enrichment of 11.74 and *adj.p* = 2.12E-52, indicating its potential involvement in cell cycle regulation and G2/M checkpoint signalling in cancer. ***Figure 3C*** shows a bubble plot visualizing the enrichment results, where bubble size indicates gene count, color represents the adjusted *p*-value, and the x-axis denotes the gene ratio.

**Table 1:** Over-representation analysis of 256 genes obtained from the intersection of differentially expressed genes (DEGs) and KNL1 associated green module.

| **Hallmark Description** | **Gene Count** | **Fold-Enrichment / p.adj** | **Gene IDs** |
|---|---|---|---|
| Hallmark E2F Targets | 60 | 11.74 / 2.12E-52 | *STMN1/DIAPH3/DLGAP5/SPC25/DEPDC1/MELK/UBE2T/KIF4A/CENPM/CDKN3/CDK1/CENPE/PCNA/E2F8/PTTG1/SPC24/BUB1B/CDKN2A/TOP2A/MCM5/MAD2L1/PRIM2/KPNA2/RNASEH2A/CDC2/ASF1B/CCNB2/MCM2/HMGA1/RRM2/RAD51AP1/KIF2C/MCM6/PLK4/HMMR/MKI67/AURKB/RACGAP1/KIF18B/TCF19/TRIP13/GINS1/CDCA8/MCM4/MYBL2/CDKN2C/BIRC5/DSCC1/TK1/AURKA/CDC25A/CCNE1/CIT/CHEK1/ORC6/UBE2S/CKS2/GINS4/CDCA3/PLK1* |
| Hallmark G2M Checkpoint | 60 | 11.74 / 2.12E-52 | *STMN1/PBK/NEK2/EXO1/NDC80/KIF4A/BUB1/CDKN3/TTK/CDK1/CENPA/CENPE/CENPF/PTTG1/CDC45/TOP2A/MCM5/MAD2L1/UBE2C/PRIM2/KPNA2/KIF15/CDC20/CCNB2/MCM2/HMGA1/KIF11/E2F1/KIF23/KIF2C/MCM6/PLK4/HMMR/MKI67/PRC1/AURKB/RACGAP1/GINS2/RAD54L/TPX2/MYBL2/CDKN2C/BIRC5/CHAF1A/TROAP/AURKA/KNL1/TRAIP/CDC25A/POLQ/E2F2/CHEK1/ORC6/UBE2S/NUSAP1/CCNF/CCNA2/CKS2/CDC6/PLK1* |
| Hallmark Mitotic Spindle | 24 | 4.72 / 7.51E-10 | *NEK2/DLGAP5/NDC80/KIF4A/BUB1/TTK/CDK1/CENPE/ANLN/CENPF/TOP2A/KIF15/CCNB2/KIF11/KIF23/KIF2C/PRC1/RACGAP1/TPX2/BIRC5/AURKA/NUSAP1/ECT2/PLK1* |
| Hallmark Spermatogenesis | 10 | 2.9 / 0.02 | *NEK2/NCAPH/BUB1/CDKN3/TTK/CDK1/CCNB2/KIF2C/AURKA/DMC1* |

As these genes belong to the same WGCNA module, we hypothesize that they exhibit coordinated expression patterns and may participate in a shared regulatory network. Their collective enrichment in the G2M checkpoint hallmark further suggests potential functional interactions in modulating G2M signalling in HCC.

#### 3.2.3. Hub Gene Identification in the G2M Checkpoint Network

A total of 60 genes were identified at the intersection of the differentially expressed genes (DEGs), the KNL1-associated green module, and the Hallmark G2M Checkpoint gene set, as illustrated by the venn diagram (***Figure 3D***).To explore the interaction landscape of these genes, a PPI network was constructed using the STRING database via the STRING plugin in Cytoscape (***Figure 3E***). The network was highly interconnected, consisting of 60 nodes and 1,708 edges, with a high average node degree (56.93), clustering coefficient (0.975), and density (0.965), indicating a compact and co-regulated structure.

To identify key regulators, nodes were ranked using Maximal Clique Centrality (MCC) via the cytoHubba plugin. The top 10 hub genes-*KNL1, CHEK1, KPNA2, EXO1, PTTG1, CCNB2, TPX2, RACGAP1, CDC20, and CDK1*-emerged as central components of the G2/M checkpoint regulatory network (***Figure 3F***).

### 3.3. Single-Sample Gene Set Enrichment Analysis (ssGSEA) Recapitulates a Tumor Subpopulation with Elevated Activity of KNL1 associated G2M network

The ten hub genes identified from the G2M checkpoint PPI network were utilized to compute a KNL1-associated G2M network score for primary tumor samples in the TCGA-LIHC cohort using single-sample Gene Set Enrichment Analysis (ssGSEA). Based on the median score, patients were stratified into High (*n* = 177) and Low (*n* = 179) score groups. A heatmap of the full 60-gene Hallmark G2M checkpoint signature revealed marked upregulation in tumor samples compared to normal liver tissues, with further elevation observed in the High-score subgroup. This suggests the existence of a tumor subpopulation characterized by heightened KNL1 associated G2M network activity **(*Figure 4A*).**

**Figure 4:**
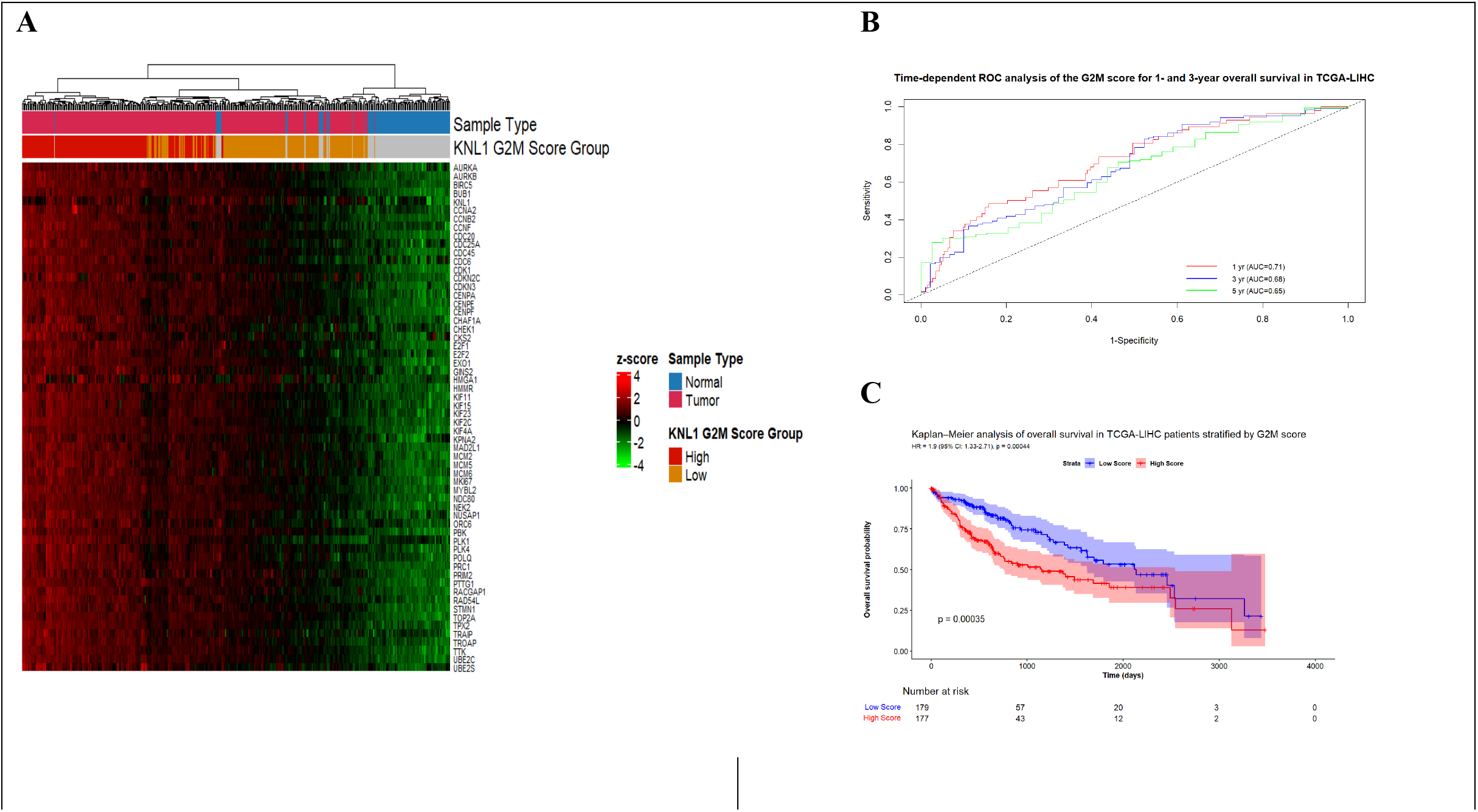
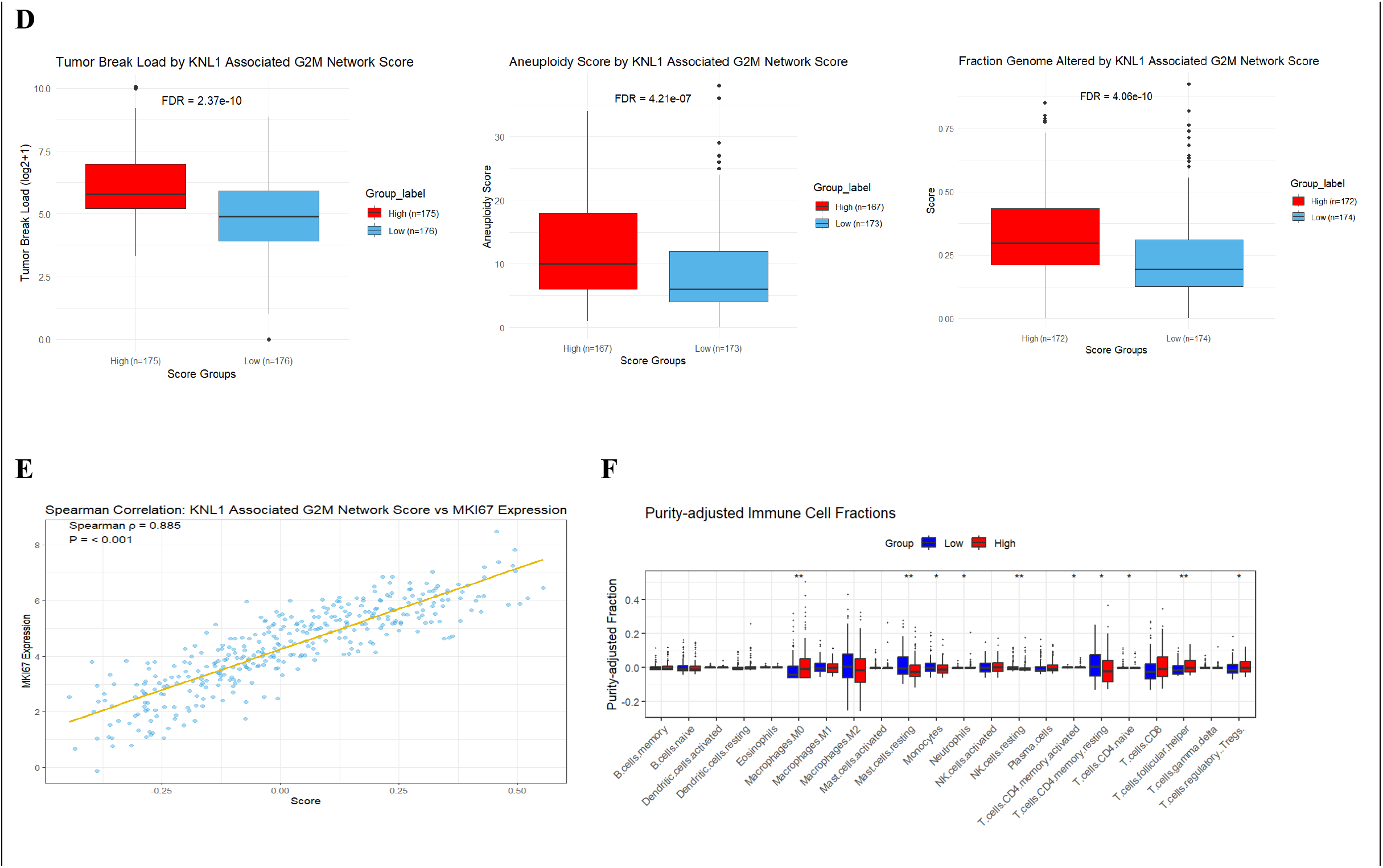
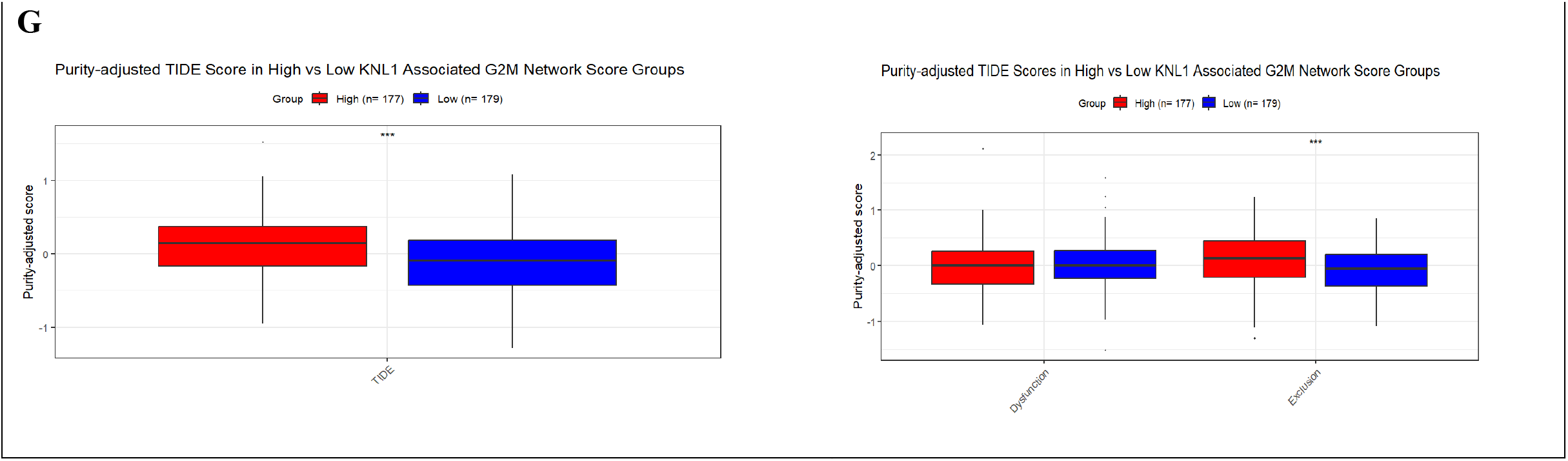
Prognostic, proliferative, genomic, and immunological implications of KNL1-associated G2M network activity in hepatocellular carcinoma. **(A)** Heatmap showing z-score–normalized expression of KNL1-associated G2M network genes across TCGA-LIHC samples, annotated by tumor versus normal status and stratified into high- and low-G2M network score groups. **(B)** Kaplan–Meier overall survival (OS) curves comparing patients with high (n = 177) and low (n = 179) G2M network scores, dichotomized by the median score in TCGA-LIHC cohort. Patients in the high-score group exhibited significantly poorer survival (HR = 1.90, 95% CI: 1.33–2.71, *p* = 4.4 × 10⁻⁴). **(C)** Time-dependent ROC analysis evaluating the prognostic performance of the G2M network score for overall survival, demonstrating moderate predictive accuracy at 1, 3, and 5 years (AUC = 0.71, 0.68, and 0.65, respectively). **(D)** Boxplots showing that tumors with high G2M network scores display significantly increased tumor break load (median [IQR]: 45 [36–126] vs. 29 [14–59.25], median Δ= 16, adjusted *p* = 2.37 × 10⁻¹⁰), aneuploidy score (10 [6–18] vs. 6 [4–12], median Δ= 4, adjusted *p* = 4.21 × 10⁻⁷), and fraction of genome altered (0.296 [0.21–0.433] vs. 0.194 [0.13–0.31], median Δ= 0.102, adjusted *p* = 4.06 × 10⁻¹⁰), indicating elevated chromosomal instability. **(E)** Spearman correlation analysis demonstrating a strong positive association between the G2M network score and *MKI67* expression (ρ = 0.885, *p* < 0.01), consistent with enhanced proliferative activity. **(F)** Tumor purity–adjusted CIBERSORTx immune deconvolution revealed increased infiltration of regulatory T cells (median Δ = 0.0158; adjusted *p* = 4.4 × 10⁻⁴), T follicular helper cells (median difference = 0.0134; FDR = 0.0013), and M0 macrophages (median Δ = 0.0319; adjusted *p* = 0.00437) in high-score tumors. **(G)** TIDE analysis indicated higher immune evasion potential in the high-score group, with significantly elevated TIDE scores (median [IQR]: 0.14 [−0.166–0.372] vs. −0.09 [−0.43–0.181], median Δ= 0.23, adjusted *p* = 1.20 × 10⁻⁵) and exclusion scores (0.14 [−0.206–0.446] vs. −0.06 [−0.368–0.201], median Δ= 0.20, adjusted *p* = 1.86 × 10⁻⁴) after adjustment for tumor purity suggesting enhanced immune evasion potential. All group comparisons were performed using the Wilcoxon rank-sum test. Data are presented as median and interquartile range (IQR). Multiple testing correction was applied using the Benjamini–Hochberg method, with adjusted *p* < 0.05 considered statistically significant.

Importantly, the expression profile of the ten selected hub genes recapitulated the broader activity pattern of the full 60-gene G2M signature, indicating that this refined gene set effectively captures the core regulatory features of the pathway. This supports the utility of the hub-gene–based network score as a surrogate for the whole 60-gene G2M signature.

#### 3.3.1 High KNL1 Associated G2M Network Scores Predict Poor Overall Survival in TCGA-LIHC Cohort

To evaluate the prognostic relevance of cell cycle activity, we analysed KNL1 G2M network score as a continuous variable. Higher G2M scores were significantly associated with worse overall survival (OS) in both univariate analysis (HR = 1.56 per SD increase, *p* = 1.5 × 10⁻⁶) and multivariable Cox regression (HR = 1.46, 95% CI: 1.09–1.95, *p* = 0.01), after adjustment for age, sex, pathologic stage, histologic grade, viral status, vascular invasion, and serum fetoprotein levels.

Among the clinical covariates, older age at diagnosis was independently associated with poorer survival (HR = 1.03 per year, 95% CI: 1.01–1.06, *p* = 0.007). Active HBV infection was also significantly associated with worse prognosis (HR = 2.04, 95% CI: 1.15–3.63, p = 0.015), whereas HCV infection alone was not statistically significant (*p* = 0.44).

Macrovascular invasion emerged as a strong adverse prognostic factor (HR = 5.11, 95% CI: 1.93–13.57, p = 0.001), while microvascular invasion was not independently associated with OS. Serum fetoprotein levels showed a statistically significant but modest association with survival (p = 0.021). In contrast, gender, histologic grade, and pathologic stage were not independently associated with OS in the adjusted model.

The model demonstrated good discriminatory performance (concordance index = 0.73) and was statistically significant overall (likelihood ratio test p = 7e-07). Time-dependent ROC curves demonstrated moderate prognostic accuracy (AUC: 0.71 at 1 year; 0.68 at 3 years; 0.65 at 5 years) **(*Figure 4B*)**. Consistently, Kaplan–Meier analysis showed that patients with high G2M scores, dichotomized by the median, experienced significantly shorter OS compared with those with low scores (HR = 1.90, 95% CI: 1.33–2.71, *p* = 4.4 × 10⁻⁴) **(*Figure 4C*)**. In contrast, KNL1 expression alone was not significantly associated with OS ***(Supplementary Fig. S3A),*** supporting the superior predictive value of pathway-level scoring.

#### 3.3.2. Validation of Prognostic Performance in Independent GEO Cohorts

In LIRI-JP cohort (n = 67; high = 33, low = 34), univariate Cox regression for both the dichotomized G2M score (HR = 9.71, 95% CI: 2.72–34.61, *p* = 4.58 × 10⁻⁴; ***Supplementary Fig. S3B***) and the continuous G2M score (HR = 3.36, 95% CI: 1.63–6.94, *p* = 0.00106) was significantly associated with worse survival. These associations remained independent predictors of mortality after adjustment for age and sex, stage and tumor grade.

In GSE14520, the G2M network score retained prognostic significance in the GSE14520 microarray cohort despite partial probe coverage for KNL1. Higher continuous G2M scores were associated with worse OS (HR = 1.39 per SD, 95% CI: 1.09–1.76, *p* = 0.007), these results remained significant after adjusting for age, sex, AFP level, and HBV infection status. Additionally, high-score patients (n= 113) showed a two-fold higher risk of death (HR = 2.05, 95% CI: 1.32–3.17, *p* = 0.001) compared with low score patients (n= 112) ***(Supplementary Fig. S3C)***. Full multivariate effect estimates are reported for TCGA-LIHC ***(Supplementary Table S1)***, LIRI-JP ***(Supplementary Table S2),*** and GSE14520 ***(Supplementary Table S3)*.**

Time-dependent ROC analysis demonstrated stable prognostic performance in the GSE14520 cohort, with AUCs of 0.60, 0.62, and 0.61 at 1-, 3-, and 5-year time points. In the LIRI-JP cohort, prognostic accuracy was evaluated at 1 and 3 years, yielding AUCs of 0.79 and 0.84, respectively; 5-year ROC analysis was not performed because all patients had experienced the event by this time point, precluding discrimination.

In the GSE54236 cohort (n = 81; high = 41, low = 40), using 1-year overall survival as a binary outcome, higher G2M activity was significantly associated with poorer survival when modeled as a continuous variable (HR = 3.48 per SD increase; 95% CI: 1.85–6.55; *p* = 1.12 × 10⁻⁴). Dichotomized analysis showed a similar direction of effect (HR = 2.97); however, statistical significance was not achieved, likely reflecting the limited number of events (n = 15).

### 3.4. High KNL1 Associated G2M Scores Correlates with Higher Expression of Proliferation Marker MKI67 and Showed Significantly Higher Chromosomal Instability Scores

Chromosomal instability and mutation analyses revealed that the high-score group exhibited significantly higher tumor break load (median [IQR]: 45 [36–126] vs. 29 [14–59.25], median Δ= 16, *adj.p* = 2.37 × 10⁻¹⁰), aneuploidy score (10 [6–18] vs. 6 [4–12], median Δ= 4, *adj.p* = 4.21 × 10⁻⁷), and fraction of genome altered (0.296 [0.21–0.433] vs. 0.194 [0.13–0.31], median Δ= 0.102, *adj.p* = 4.06 × 10⁻¹⁰) compared to the low-score group. However, no significant difference was detected in the tumor mutation burden (median [IQR]: 2.92 [2.14–4.09] vs. 2.87 [2.03–4.00], median Δ= 0.05, *adj.p* = 0.785; ***Supplementary Figure S4***). These findings suggest that the identified KNL1 associated G2M network gene signature may contribute to the promotion of chromosomal instability in HCC, a well-established hallmark of cancer (102) **(*Figure 4D*)**.

This distinction is biologically plausible, as TMB primarily reflects the accumulation of point mutations and small insertions or deletions, often driven by defects in DNA mismatch repair pathways (34,35,103). In contrast, large-scale chromosomal alterations and structural variations—key features of chromosomal instability—are more accurately captured by metrics such as tumor break load, aneuploidy, and FGA, underscoring the limited utility of TMB for characterizing chromosomal instability in HCC (103).

Additionally, to further evaluate the proliferative status of tumors in the high-score group, we performed a Spearman correlation analysis between the KNL1-associated G2M Network score and MKI67 expression. The analysis revealed a strong positive correlation (ρ = 0.885, *p* < 0.01), indicating that tumors with high G2M scores exhibit enhanced proliferative capacity **(*Figure 4E*)**.

### 3.5. High KNL1-G2M Network Activity Correlates with Differential Immune Infiltration and Higher TIDE Exclusion Score

To characterize immune microenvironment differences associated with KNL1–G2M pathway activation, we compared immune cell abundances after purity adjustment between the high-and low-G2M groups. Several adaptive immune cell subsets were significantly enriched in the high-score group, including T follicular helper cells (median Δ= 0.0134; *adj.p* = 0.0013) and regulatory T cells (Tregs; median Δ= 0.0158; *adj.p* = 0.00044). In contrast, T cells CD4 memory resting were markedly reduced (median Δ= −0.0294; *adj.p* = 0.0123).

Innate immune populations also demonstrated substantial shifts. M0 macrophages were increased in the high group (median Δ= 0.0319; *adj.p* = 0.00437), whereas resting NK cells (median Δ= −0.0044; *adj.p* = 0.00437) and resting mast cells (median Δ= −0.0203; *adj.p* = 0.00437) were significantly reduced **(*Figure 4F*)**.

Given that the TIDE scoring system was primarily developed using cancers other than HCC, its predictions should be interpreted cautiously (83). Nevertheless, consistent with this altered immune landscape, tumors in the high-G2M group demonstrated greater immune evasion potential based on TIDE predictions. High-score tumors exhibited significantly elevated overall TIDE scores (median [IQR]: 0.14 [−0.166–0.372] vs. −0.09 [−0.43–0.181], median Δ= 0.23, *adj.p* = 1.20 × 10⁻⁵) and a markedly higher exclusion score (0.14 [−0.206–0.446] vs. −0.06 [−0.368–0.201], median Δ= 0.20, *adj.* = 1.86 × 10⁻⁴), indicating impaired T-cell infiltration and a microenvironment more permissive to immune escape **(*Figure 4G*)**. These results suggest that tumors with heightened G2M pathway activity—are more capable of evading immune surveillance.

### 3.6. Single-Cell Analysis Identifies a Proliferative Hepatocyte Subpopulation With High KNL1-Associated G2M Network Activity

After quality control, 62,537 high-quality cells from the GSE149614 dataset were retained, spanning primary tumors, portal vein tumor thrombus (PVTT), metastatic lymph nodes, and non-tumor liver tissues. Batch correction and integrated clustering resolved 16 transcriptionally distinct cell populations, including hepatocytes, endothelial cells, T/NK cells, B cells, myeloid cells, and fibroblasts. Hepatocyte-enriched clusters were primarily represented by clusters 0, 8, and 15, as shown in the UMAP and cell type annotations **(*Figure 5A-B*).**

**Figure 5:**
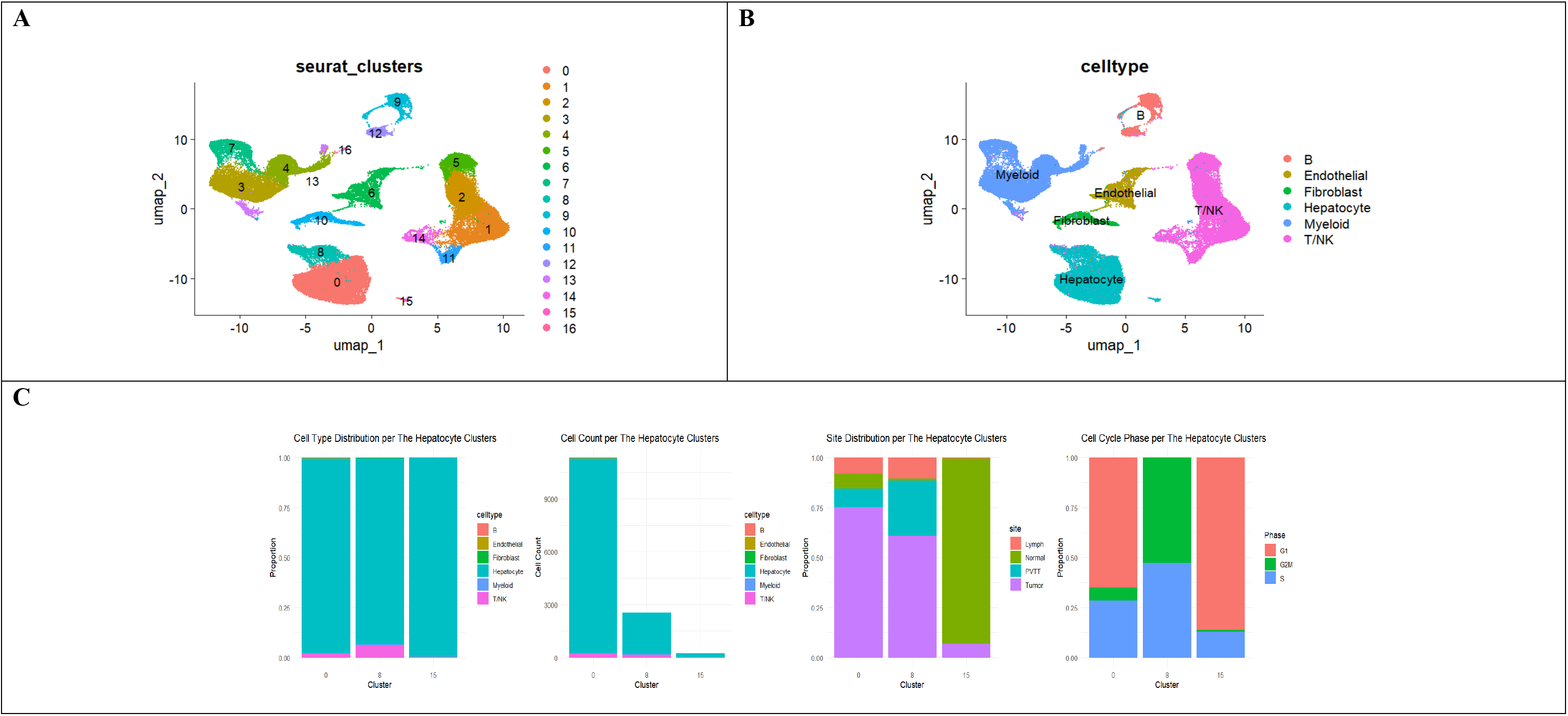
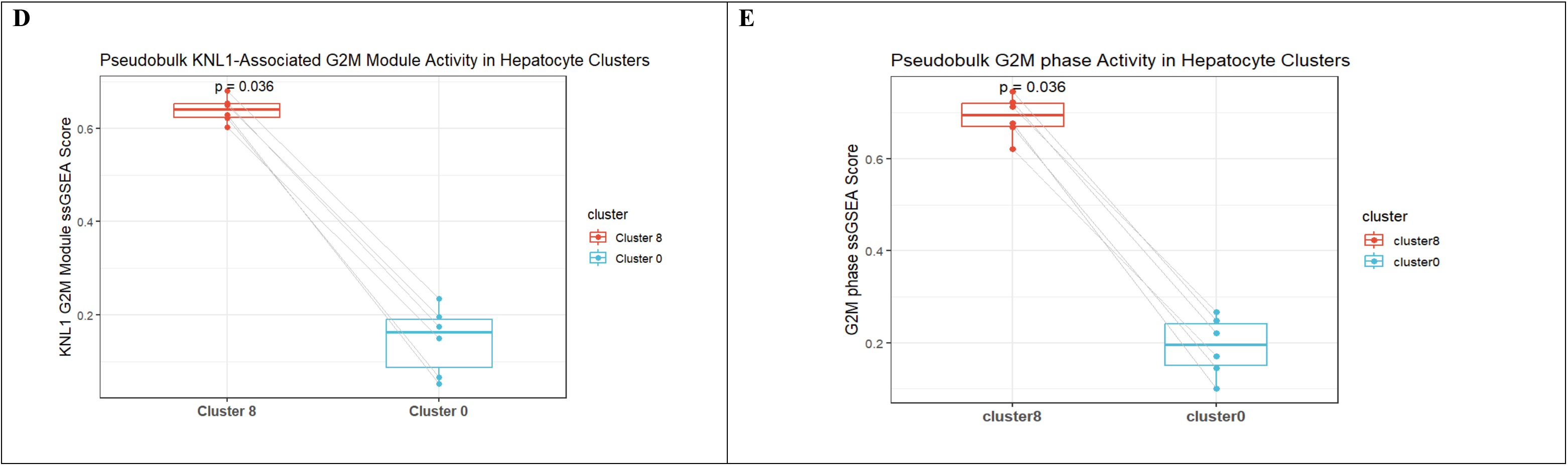
Single-cell characterization of hepatocyte clusters and KNL1-associated G2M network activity in the GSE149614 dataset. **(A)** UMAP visualization of 62,537 high-quality cells resolved into 16 transcriptionally distinct clusters following quality control, batch correction, and integration. **(B)** Cell type annotation identifying major populations, including hepatocytes, endothelial cells, T/NK cells, B cells, myeloid cells, and fibroblasts. **(C)** Bar plots summarizing features of hepatocyte-enriched clusters (0, 8, and 15), including cell type composition, total cell number, tissue site distribution, and cell cycle phase proportions. Cluster 8 is enriched for tumor and PVTT-derived cells and exhibits increased G2/M and S phase fractions. **(D)** Boxplot showing pseudobulk KNL1-associated G2/M network activity in hepatocyte clusters 8 and 0. Scores were computed by ssGSEA, revealing significantly higher KNL1-associated G2/M activity in cluster 8 (paired Wilcoxon test, p = 0.031; median Δ = +0.47). **(E)** Boxplot showing pseudobulk G2/M phase activity in hepatocyte clusters 8 and 0. G2/M activity scores were calculated by ssGSEA using the Seurat cc.genes.updated gene set on pseudobulked single-cell RNA-seq data. Cluster 8 shows significantly higher G2/M activity (paired Wilcoxon test, p = 0.031; median Δ = +0.50). For panels **(D–E),** each point represents a pseudobulk sample, with lines connecting matched samples across clusters. Boxes indicate median and interquartile range (IQR).

Further characterization of the hepatocyte clusters is shown in ***Figure 5C***. All three clusters—0, 8, and 15—were composed almost exclusively of hepatocytes. Cluster 0 contained the majority of hepatocytes and was largely derived from tumor tissues. Cluster 8 included hepatocytes primarily from tumor samples and portal vein tumor thrombus (PVTT), while cluster 15 was predominantly composed of hepatocytes from non-tumor liver tissues.

Cell-cycle phase analysis highlighted prominent differences between hepatocyte subsets. Cluster 8 displayed strong enrichment for G2/M and S phase cells, with almost undetectable G1 cells, indicating highly active proliferation. In contrast, cluster 0 exhibited a more balanced distribution dominated by G1, whereas cluster 15 displayed a cell-cycle profile consistent with non-malignant hepatocytes.

### 3.7. Pseudobulk Analysis Confirms Cluster-Specific Proliferative Signatures

To ensure that observed differences reflect true patient-level biology, pseudobulk aggregation was performed for each patient × cluster combination. After applying thresholds of ≥100 cells and ≥10⁴ UMIs, six patients remained for paired analyses between tumor-derived hepatocyte clusters.

Across all matched patients, cluster 8 consistently exhibited markedly higher KNL1-associated G2M module activity compared with cluster 0 (paired Wilcoxon p = 0.031; rank-biserial r = 1.00; median Δ = +0.47) **(*Figure 5D*)**. G2/M-phase pathway activity followed the same pattern, with cluster 8 showing significantly elevated cell-cycle progression relative to cluster 0 (paired Wilcoxon p = 0.031; rank-biserial r = 1.00; median Δ = +0.50) **(*Figure 5E*)**. These effect sizes indicate a uniform proliferative shift across patients.

Pseudobulk differential expression further validated these observations, with KNL1 significantly upregulated in cluster 8 versus cluster 0 (log₂FC = 1.59; *adj.p* = 0.0013). This transcriptional increase aligns with the elevated G2M pathway activity and supports the identification of cluster 8 as a high-mitotic, KNL1-high malignant hepatocyte subpopulation. Cluster 15, representing predominantly normal hepatocytes, did not meet the ≥100-cell criterion in most samples and was therefore excluded from pseudobulk testing.

### 3.8. KNL1-High Tumor Cells Exhibit Enhanced ECM and Immune Signaling

To investigate how KNL1 expression impacts cell–cell communication, we used CellChat to compare tumor cells with High KNL1 associated G2M network module score and positive KNL1 expression (High_Positive, *n= 1046*) versus those with Low scores and undetectable KNL1 expression (Low_Negative, *n= 13680*). Overall, the High_Positive group displayed a markedly increased number and strength of inferred cell–cell interactions (***Figure 6A*)**. Heatmap visualization revealed increased incoming signals from fibroblasts to hepatocytes in the High_Positive group, suggesting enhanced crosstalk within the tumor microenvironment (***Figure 6B*)**. Information flow analysis showed greater signal transmission through both ECM and immune-related pathways in the High_Positive condition (***Figure 6C*)**.

**Figure 6:**
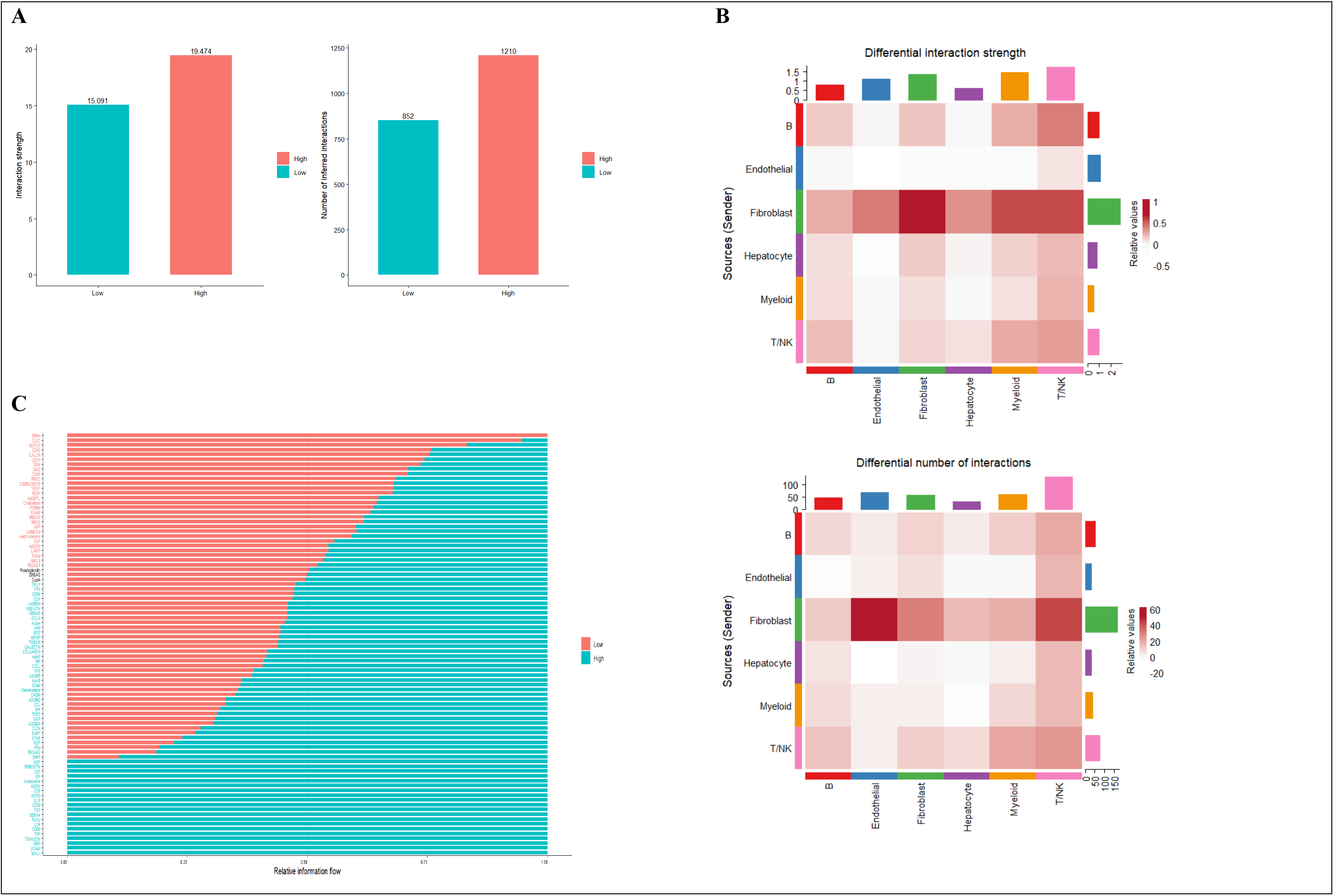
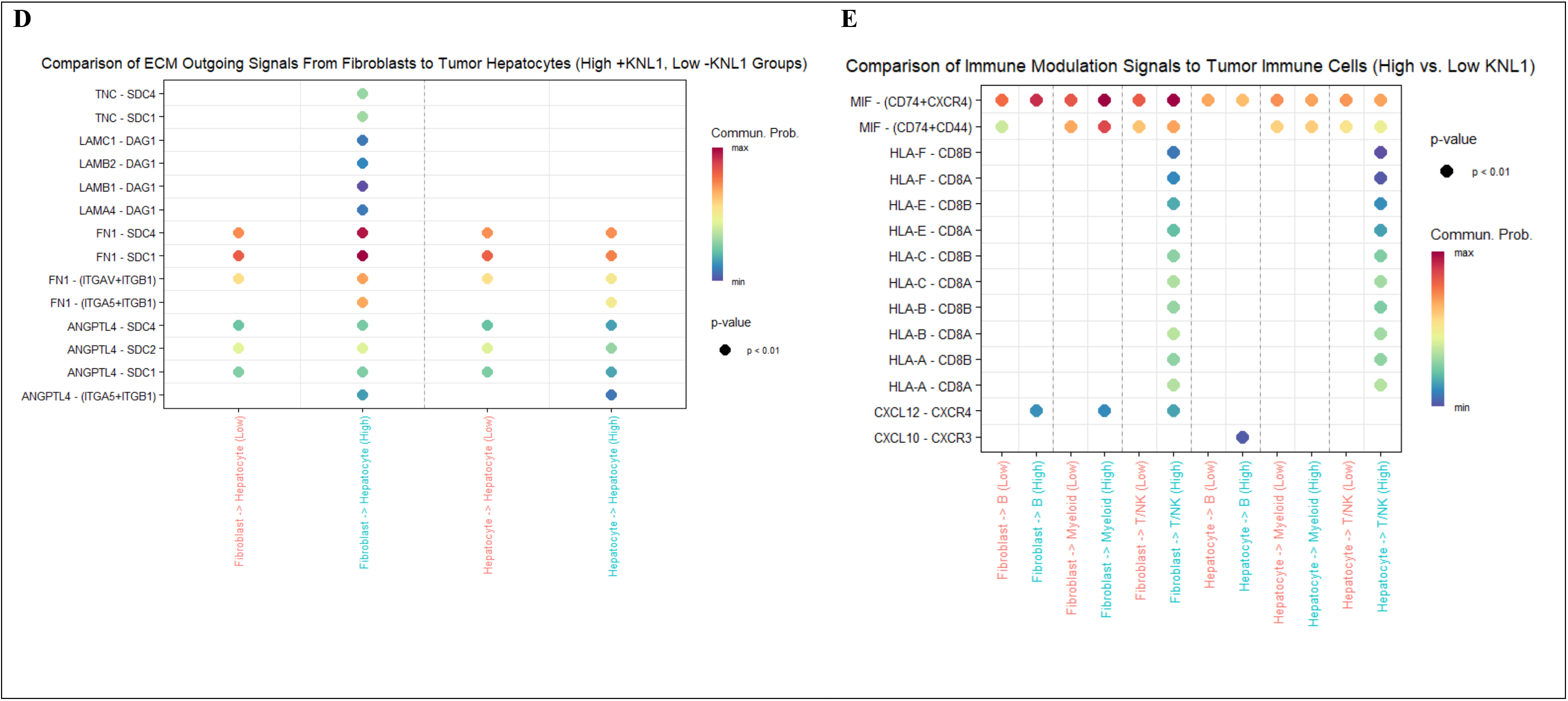
Cell–cell communication analysis between High KNL1-associated G2M module score and KNL1-positive tumor cells (High_Positive) and Low KNL1-associated G2M score and KNL1-negative tumor cells (Low_Negative). **(A)** CellChat differential interaction bar plot showing increased number and strength of interactions in the High_Positive (n= 1046) compared to Low_Negative (n= 13680) tumor cells. **(B)** CellChat differential interaction heatmap showing increased incoming signalling to hepatocytes from fibroblasts in the High_Positive relative to low_negative tumor cells, indicating enhanced tumor–stromal crosstalk. **(C)** Information flow analysis revealing elevated signalling strength in High_Positive tumors across extracellular matrix and immune-related pathways. **(D)** Bubble plot illustrating enriched ECM-derived signals to High_Positive hepatocytes via ligands such as tenascins (TNC), laminins (LAMC1, LAMB2, LAMB1, LAMA4), fibronectin 1 (FN1), angiopoietin like protein 4 (ANGPTL4), targeting integrin, syndecan, and dystroglycan 1 (DAG1) receptors. **(E)** Bubble plot showing enhanced immune signalling in High_Positive tumors, including MHC-I signalling (HLA-A/B/C/E/F → CD8A/B) and CXCL10–CXCR3 signalling from hepatocytes to B cells. In bubble plots, circle size represents communication probability, and color intensity reflects statistical significance (p-value) as computed by CellChat. [TNC, tenascin C; LAMC1, Laminin subunit gamma-1; LAMB2, Laminin subunit beta-2; LAMA4, Laminin Subunit Alpha 4, ANGPTL4, angiopoietin like protein 4; DAG1, dystroglycan 1; MHC-I, major histocompatibility complex class I; HLA, human leukocyte antigen; CD8, cluster of differentiation 8; CXCL10, C-X-C motif chemokine ligand 10, CXCR3, C-X-C motif chemokine receptor 3].

#### 3.8.1. ECM–Tumor Crosstalk Mediated by Integrins and Syndecans

To further characterize ECM-related communication directed toward tumor hepatocytes, we examined incoming ECM signaling at the ligand–receptor interaction level. Bubble plot analysis revealed that High_Positive hepatocytes received significantly more signals from fibroblasts via ECM ligands such as tenascins (TNC), laminins (LAMC1, LAMB1, LAMB2, and LAMA4), fibronectin (FN1), angiopoietin-like protein 4 (ANGPTL4). These signals predominantly targeted integrin, syndecan, and dystroglycan 1 (DAG1) receptors enriched in High_Positive cells **(*Figure 6D*)**.

#### 3.8.2. Immune Signalling Differences

To explore immune signaling within the tumor microenvironment in High_Positive vs Low_Negative cells, we assessed intercellular communication across three key immune-related pathways: MIF, MHC-I and CXCL, based on the previous information flow analysis. Bubble plot analysis revealed significantly enriched MHC-I signaling in the High_Positive tumors, characterized by enhanced engagement of HLA class I molecules (HLA-A/B/C/E/F) from fibroblasts, and hepatocytes toward CD8A/CD8B receptors on T and NK cells. Notably, CXCL signalling was also enhanced, particularly involving CXCL10 secreted by tumor hepatocytes and CXCR3 receptors on B cells **(*Figure 6E*).**

Collectively, these findings indicate that the KNL1-associated G2M network is linked to coordinated remodeling of extracellular matrix–dependent communication and immune signaling within the hepatocellular carcinoma microenvironment.

### 3.9. KNL1 was successfully downregulated in HepG2 Cells

To assess the efficiency of KNL1 knockdown in HepG2 cells, we first evaluated transfection success based on GFP reporter expression. Mean fluorescence intensity (MFI) quantified via ImageJ showed a substantial increase in the KNL1 gRNA1+2 group, with an approximately 11-fold elevation compared to control cells (22.9 ± 1.59 vs. 2.05 ± 0.068, *p= 0.001*) **(*Figure 7A*).**

**Figure 7:**
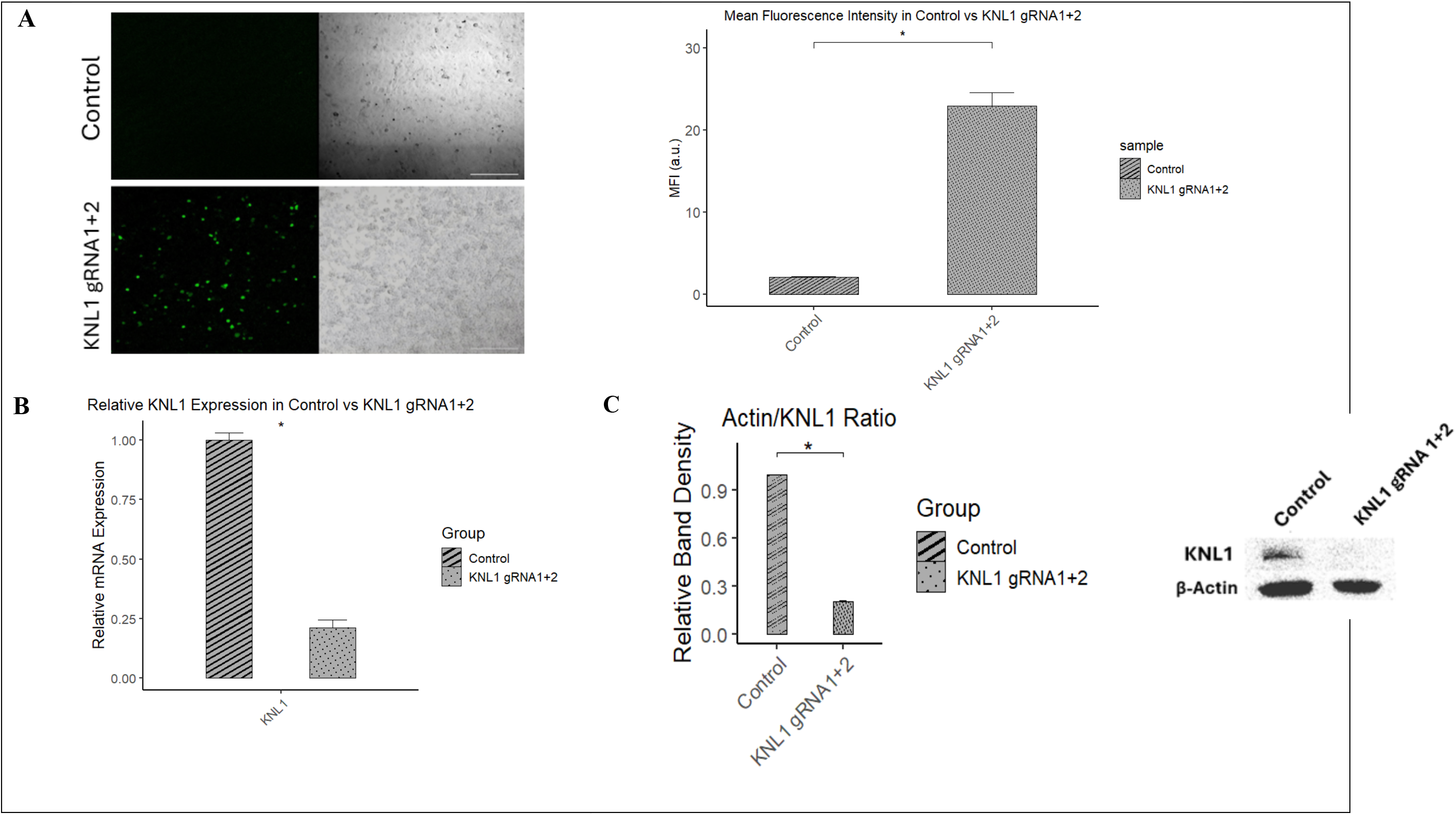
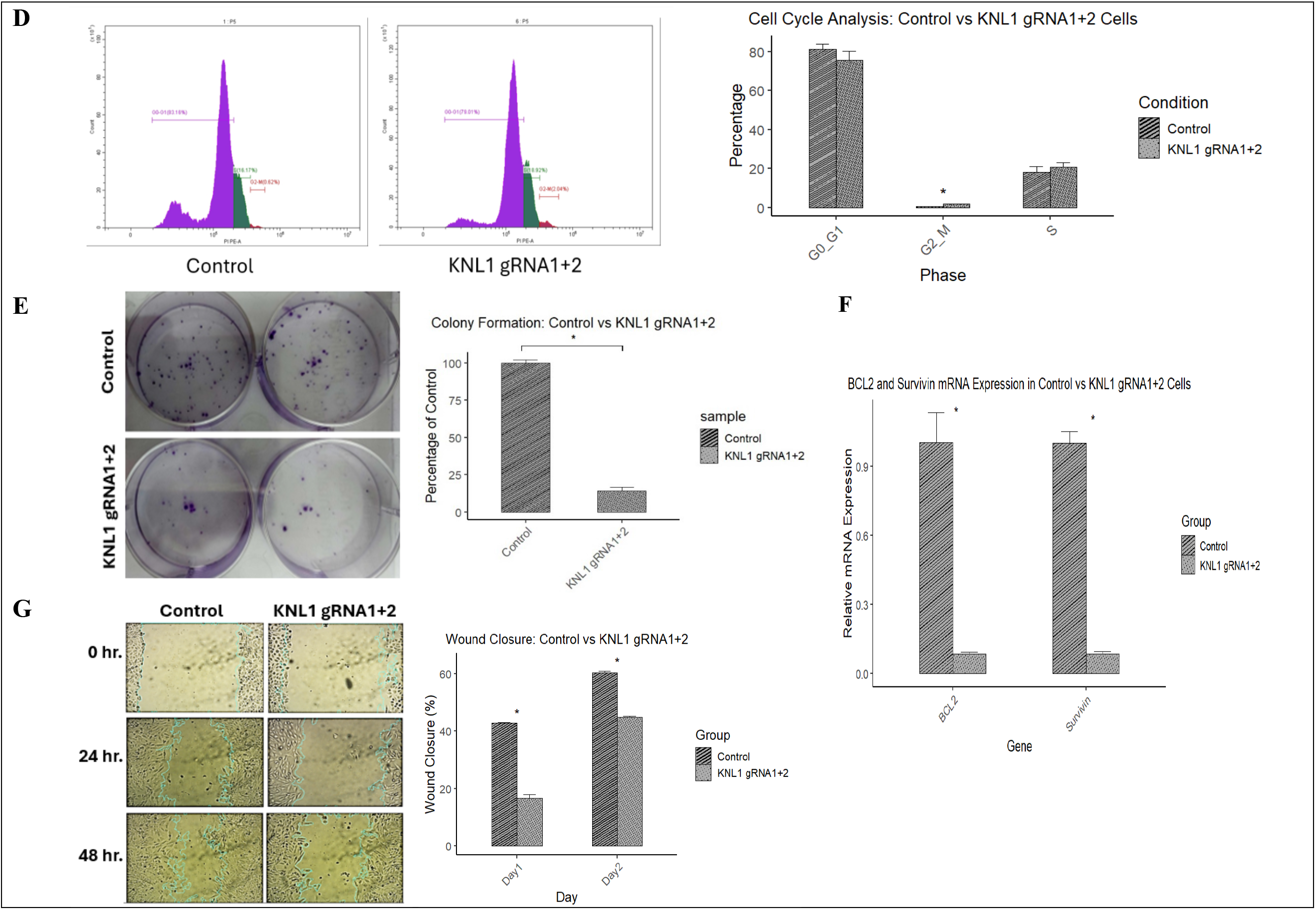
KNL1 downregulation in HepG2 cells reduces proliferation, induces G2/M arrest and downregulated migration of HepG2. **(A)** Representative confocal images showing GFP expression 72 h post-transfection with KNL1-targeting vectors (left) and quantification of mean fluorescence intensity (MFI; right), confirming successful transfection. Scale bar: 200 μm. (B) RT-qPCR analysis demonstrating significant downregulation of KNL1 mRNA in KNL1 gRNA1+2–treated cells compared with control (0.212 ± 0.03 vs. 1.00 ± 0.03; p = 0.002). Relative expression levels were calculated using the 2⁻ΔΔCt method, normalized to GAPDH, and expressed as fold change relative to control. (C) Representative Western blot images and densitometric quantification of KNL1 protein levels normalized to β-actin. (D) Flow cytometry analysis showing a significant increase in the proportion of G2/M-phase cells following KNL1 downregulation (1.97 ± 0.09% vs. 0.635 ± 0.021%; p = 0.027). (E) Colony formation assay showing representative images after 10 days and quantification of colony numbers, revealing a marked reduction in clonogenic capacity in KNL1 gRNA1+2–treated cells compared with control (20 ± 4 vs. 138 ± 3 colonies; p = 0.001). (F) RT-qPCR analysis demonstrating significant downregulation of BIRC5 (Survivin; 0.09 ± 0.01 vs. 1.00 ± 0.05) and BCL2 (0.085 ± 0.008 vs. 1.00 ± 0.128) in KNL1 gRNA1+2–treated cells. Relative expression levels were calculated using the 2⁻ΔΔCt method and normalized to a housekeeping gene. (G) Wound healing assay showing representative images (10× magnification) at 0, 24, and 48 h post-scratch (left) and quantification of wound closure (right). KNL1 downregulation significantly reduced migration at 24 h (42.8% ± 0.35 vs. 33.6% ± 1.27; p = 0.001) and 48 h (60.4% ± 0.59 vs. 44.8% ± 0.35; p = 0.001).

RT-qPCR analysis confirmed effective KNL1 downregulation at the transcript level, revealing a 4.7-fold reduction in mRNA expression in the KNL1 gRNA1+2 cells relative to control (0.212 ± 0.0315 vs. 1.00 ± 0.0307; *p* =0.002) **(*Figure 7B*).**

Consistently, Western blot analysis demonstrated significant downregulation of KNL1 protein following CRISPR-Cas9 treatment, showing an approximate 4.8-fold decrease in expression normalized to β-actin, compared to control (*p* = 0.003) **(*Figure 7C*).**

### 3.10. KNL1 downregulation in HepG2 cells induces G2/M arrest, reduces proliferation, and suppresses the migration

KNL1 downregulation led to a significant accumulation of cells in the G2/M phase, with a 3.1-fold increase in G2/M-phase cells compared to the control group (1.97 ± 0.09% vs. 0.635 ± 0.021%; *p* = 0.027) **(*Figure 7D*)**. Proliferation capacity was markedly impaired, as shown by an ∼85% reduction in colony formation in the KNL1 gRNA1+2 cells compared to controls (138 ± 3 vs. 20 ± 4 colonies*, p* = 0.001) **(*Figure 7E*).** This was accompanied by a reduction in the expression of the anti-apoptic markers Bcl-2 (0.085<u>1</u> ± 0.008 vs 1 ± 0.128) and survivin (0.09 ± 0.01 vs 1 ± 0.05) **(*Figure 7F*).**

Finally, wound healing assay demonstrated impaired migratory ability in KNL1-downregulated cells, with a 21.5% reduction in wound closure at 24 h (Control: 42.8% ± 0.345 vs. KNL1 gRNA1+2 33.6% ± 1.27, *p* = 0.001) and a 25.8% reduction at 48 h (Control: 60.4% ± 0.594 vs. KNL1 gRNA1+2 44.8% ± 0.350, *p* = 0.001) **(*Figure 7G*).**

### 3.11. KNL1 Knockdown Induces Ultrastructural Features of Anoikis

To investigate the ultrastructural effects of KNL1 knockdown, transmission electron microscopy (TEM) was performed on the control and KNL1 gRNA1+2 co-transfected HepG2 cells. As shown in ***Figure 8***, control cells **(*Figure 8A–C*)** exhibited typical HepG2 morphology, including intact plasma membranes, euchromatic nuclei with well-defined nuclear envelopes and prominent nucleoli, and clearly preserved intercellular junctions.

**Figure 8:**
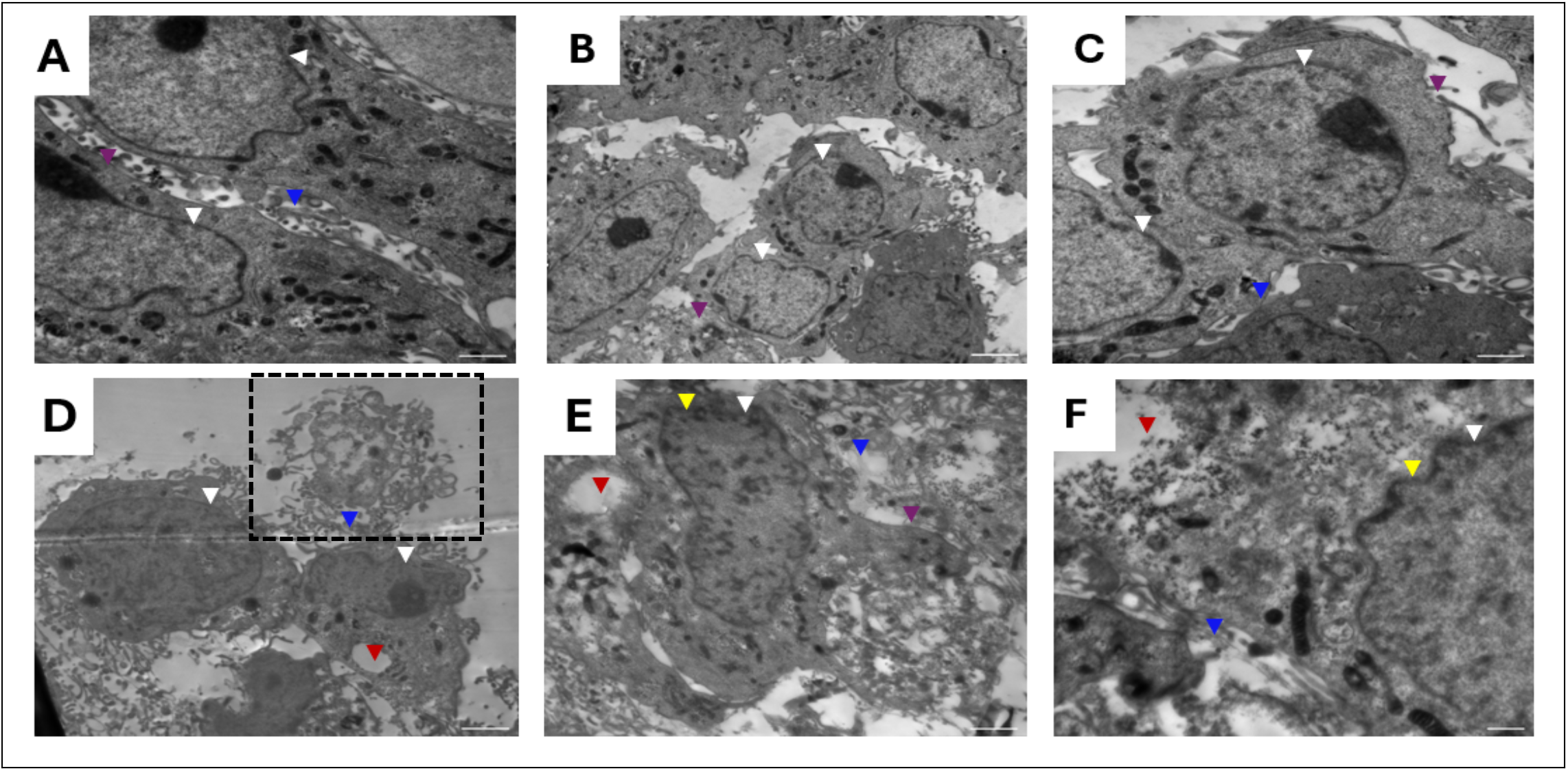
Transmission electron microscopy (TEM) reveals ultrastructural changes in HepG2 cells following KNL1 knockdown. Representative TEM images of control (A–C) and KNL1 gRNA1+2 (D–F) HepG2 cells. Scale bars: 2 µm (A, B, D, E, F) and 500 nm (C); magnifications: 1200× (A, E), 2000× (B), 2500× (D), 3000× (F), and 5000× (C). Color-coded arrowheads indicate key features: white, nuclear membrane; yellow, chromatin condensation; red, cytoplasmic vacuolization; purple, cell membrane.

In contrast, the treatment cells **(*Figure 8D–F*)** displayed multiple ultrastructural features suggestive of early apoptosis and potentially detachment-induced cell death (anoikis), such as irregular or blebbed plasma membranes **(*Figure 8E*),** nuclear membrane irregularities **(*Figure 8D-F*)**, chromatin condensation **(*Figure 8E-F*)**, and extensive cytoplasmic vacuolization **(*Figure 8D-F*)**.

A particularly notable observation in the KNL1 gRNA1+2 group was the disrupted cell–cell adhesion **(*Figure 8D-F*)**. In ***Figure 8D***, a detached cell displaying nuclear fragmentation and apoptotic morphology was identified, suggestive of anoikis induction following KNL1 knockdown (104).

## 4. Discussion

HCC is characterized by profound intratumoral heterogeneity shaped by genomic instability, and tumor–microenvironmental interactions (19,25). These features can collectively contribute to immune evasion, therapeutic resistance, and recurrence (19,103). Aberrant expression of kinetochore proteins disrupts chromosome alignment and segregation, leading to aneuploidy, chromothripsis, and complex structural rearrangements (38,105,106). KNL1 is well established as a central regulator of mitosis and the spindle assembly checkpoint; however, its broader molecular networks and functional implications in cancer-particularly HCC-remain insufficiently characterized. The strength of our study lies in its integrative approach, combining bulk RNA-sequencing data from TCGA-LIHC with single-cell transcriptomic profiling from GSE149614, and further substantiated by in vitro validation experiments. This convergence of multi-level analyses provides a more comprehensive understanding of KNL1’s role in HCC and offers mechanistic insights that extend beyond its canonical mitotic functions.

**First**, we found that KNL1 is significantly overexpressed in HCC tumors compared with normal liver tissue, with even higher levels in tumors harbouring TP53 mutations. To our knowledge, this is the first evidence identifying tumor break load as the principal chromosomal instability metric associated with KNL1 in HCC. Tumor break load is a recently introduced chromosomal instability metric that measures the burden of unbalanced structural variants involving DNA breaks ≥50 bp (37). Supporting its role in chromosomal instability, a pan-cancer analysis demonstrated positive correlations between KNL1 expression and chromosomal instability across multiple cancer types (40).

This preferential association with structural CIN likely arises from the molecular functions of KNL1. As a kinetochore scaffold, KNL1 recruits spindle assembly checkpoint (SAC) proteins such as BUB1, BUBR1, and MAD2 to ensure proper tension sensing and mitotic timing (107). When overexpressed or dysregulated, KNL1 weakens SAC fidelity, allowing cells with misattached or incompletely replicated chromosomes to prematurely enter anaphase. This can result in chromosome bridges, lagging chromatin, and replication stress-induced double-strand breaks, which subsequently trigger breakage–fusion–bridge cycles and large-scale chromosomal rearrangements (108). In contrast, classical numerical CIN is mainly driven by defects in centrosome amplification, spindle polarity, or cohesion loss, which directly cause whole-chromosome mis-segregation and (109). As KNL1 dysfunction mainly affects checkpoint surveillance and chromosome integrity, its downstream consequences appear to be more closely associated with structural genomic alterations rather than whole-chromosome mis-segregation.

**Second**, **to explore the KNL1-associated transcriptional network and its link to aggressive tumor phenotypes**, we applied WGCNA and identified a KNL1-associated G2M network at which KNL1 ranked among the top 10 hub genes based on maximal clique centrality. Four of these hub genes -CDC20, CDK1, CCNB2 and CHEK1-are core regulators of the spindle assembly checkpoint and DNA damage signalling that functionally intersect with KNL1 (39,110,110–115). All four are significantly upregulated in HCC and associated with poor patient prognosis (110,111,113,116). These findings highlight the strength of our integrative approach in revealing a functionally coherent, oncogenically relevant gene network centred on KNL1.

Single-sample GSEA revealed that high KNL1-network activity defines an aggressive HCC subgroup with poorer survival and elevated MKI67 expression, a finding validated by scRNA-seq showing that the KNL1-associated G2M signature is concentrated in proliferative tumor hepatocytes enriched in the G2/M phase and positively correlated with cell cycle progression. Consistent with these observations, CRISPR-Cas9-mediated dual targeting of KNL1 in HepG2 cells significantly impaired proliferation and induced G2/M arrest, directly demonstrating KNL1’s role in sustaining mitotic progression. Supporting this, a recent study reported that kaempferol treatment inhibited proliferation and triggered G2/M arrest in HCC cells through regulation of the ATM/CHEK2/KNL1 pathway, highlighting KNL1 as a potential therapeutic vulnerability in HCC (117). Similarly, KNL1 downregulation has been shown to suppress proliferation in cervical (HeLa), colorectal, endometrial, and non-small cell lung cancer cells, further reinforcing its role in promoting tumor cell proliferation (118,119,46,120).

**Third, given the established association between KNL1 and immune-evasive phenotypes,** we evaluated immune infiltration patterns in tumors with high KNL1-associated G2M network scores. These tumors displayed reduced infiltration of naïve B cells and monocytes, alongside enrichment of regulatory T cells, T-follicular helper cells and a marked increase in M0 macrophages. While M1 and M2 macrophage levels did not differ significantly between groups. The enrichment of M0 macrophages—typically regarded as undifferentiated or transitional—suggests the presence of a macrophage population that does not conform to the classical inflammatory (M1) or anti-inflammatory (M2) states (121). This observation aligns with the macrophage spectrum model proposed by Xue et al. (2014), which describes macrophage activation as a dynamic, stimulus-dependent continuum shaped by the tumor microenvironment (122).

This atypical immune composition, together with markedly elevated TIDE and exclusion scores, suggests that high KNL1-associated G2M network activity may foster a heightened capacity for immune evasion (83). Concomitantly, these tumors exhibited hallmarks of pronounced chromosomal instability alongside elevated proliferative activity. This convergence of immune suppression and genomic instability is consistent with previous reports showing that tumors with high tumor break load often co-activate cell cycle programs while repressing immune-related pathways, thereby gaining a proliferative advantage and evading immune surveillance (103).

Single-cell level cell–cell communication analysis further suggested that HCC tumor cells with high KNL1-associated G2M module scores and positive KNL1 expression (High_Positive) may contribute to ECM remodelling that favors an immune-exclusion phenotype. Such remodelling could plausibly involve both physical and biochemical mechanisms, including fibronectin/tenascin-C–mediated matrix stiffening, a process that creates dense structural barriers to immune infiltration (123–125).

Furthermore, consistent with immune-suppressive remodelling, immune-related pathways were also enriched, including non-classical MHC-I molecules (HLA-F, HLA-E) linked to NK and CD8⁺ T-cell inhibition (126–129), and enhanced CXCL10–CXCR3 signalling, which has been associated with recruitment of exhausted or regulatory T-cell subsets and promotion of angiogenesis in HCC (130). Collectively, these data delineate a subset of HCC associated with elevated chromosomal instability, increased proliferative activity, and enhanced ECM-related and immune signaling within the tumor microenvironment. These features may be relevant for understanding variability in immune responses and could inform future efforts to refine therapeutic stratification.

**Fourth, given the emerging recognition of extracellular matrix (ECM) remodelling as a form of “architectural reprogramming” of the tumor niche** (23), our analysis revealed that High_Positive tumor cells actively remodel the ECM through fibronectin, laminins, tenascin-C, angiopoietin-like protein 4, signalling primarily via integrins, syndecans, and dystroglycan 1 receptors (131). Mechanistically, these ligand–receptor axes are known to activate canonical pro-metastatic pathways—particularly focal adhesion kinase (FAK)/Src and PI3K/AKT—thereby supporting anoikis resistance, anchorage-independent survival, and invasive dissemination (132–135,135,136). Prior studies in HCC have linked integrins (ITGB1, ITGA5, ITGAV) to sorafenib resistance, vasculogenic mimicry, epithelial–mesenchymal transition (EMT), and poor prognosis (137–139).

Similarly, tenascin-C has been associated with aggressive tumor phenotypes and pro-inflammatory, pro-migratory signaling (140). Syndecans (SDC1, SDC4) can cooperate with integrins to regulate focal adhesion dynamics, facilitate invasion, and mediate chemoresistance via PI3K/AKT activation (141–143).

In line with these findings, KNL1 knockdown in HepG2 cells reduced cell motility, induced early apoptotic features suggestive of anoikis, and suppressed the expression of the anti-apoptotic markers Bcl2 and survivin.

**Regarding therapeutic implications,** our analysis positions KNL1 and its associated G2M network within a tumor phenotype characterized by heightened proliferative activity, chromosomal instability, and altered ECM and immune signalling. Collectively, our results indicate that KNL1 may represent a candidate for further investigation in aggressive HCC and provide a framework for future mechanistic and translational studies.

**Several limitations should be acknowledged**. Functional validation was performed in a single HCC cell line (HepG2), without stable knockout or overexpression/rescue experiments, and in vivo validation is lacking. Migration assays were conducted without mitomycin-C, so proliferation may have partially contributed to observed effects, and apoptosis evidence was limited to Bcl2/Survivin reduction and EM imaging, highlighting the need for quantitative confirmation. Predicted roles in ECM and immune modulation also remain to be experimentally verified and potential toxicity associated with targeting KNL1 has not been assessed. Therefore, Future studies employing stable genetic models, co-culture systems, in vivo approaches, and rescue experiments will be required to further assess the mechanistic role and therapeutic relevance of KNL1 in hepatocellular carcinoma.

## 5. Conclusion and Future Prospectives

In this study, we identified KNL1 and its associated G2M checkpoint gene signature as potential drivers of genomic instability, enhanced proliferative signaling, and immune evasion in HCC. Notably, KNL1 overexpression correlated with elevated tumor break load, implicating its potential role in driving structural variant–associated chromosomal instability. These findings position KNL1 and its gene network as promising biomarkers of aggressive tumor behavior and potential therapeutic targets in HCC.

## Data availability

The datasets analyzed in this study are publicly available. Bulk RNA-seq data for the TCGA-LIHC cohort and normal liver tissues from GTEx were obtained from the UCSC Xena Toil Recompute Compendium. Additional genomic and clinical features were retrieved from cBioPortal (TCGA PanCancer Atlas dataset). External validation cohorts included the LIRI-JP liver cancer dataset from the International Cancer Genome Consortium data portal and the microarray datasets GSE54236 and GSE14520 available from the Gene Expression Omnibus. Detailed information regarding dataset access and preprocessing is provided in the Methods section. All other data generated or analyzed during this study are included in this article and its supplementary information files.

## Author Contributions

Nermin M. Mohamed and Rania Hassan Mohamed contributed equally to this work.

Nadia M. Hamdy and Mahmoud M. Elhefnawi jointly supervised the study and shared last authorship.

Conceptualization was carried out by N.M. Hamdy and M.M. Elhefnawi.

Study design was performed by N.M. Mohamed, R.H. Mohamed, N.M. Hamdy, and M.M. Elhefnawi.

Funding acquisition was carried out by N.M. Hamdy, M.M. Elhefnawi.

In-silico analysis was conducted by N.M. Mohamed and N.M. Hamdy, while experimental validation was performed by N.M. Mohamed and R.H. Mohamed.

Data curation and interpretation were undertaken by N.M. Mohamed, R.H. Mohamed, M.M. Elhefnawi, and N.M. Hamdy.

Supervision was provided by N.M. Hamdy and M.M. Elhefnawi.

The manuscript was drafted by N.M. Mohamed and critically reviewed and edited by R.H. Mohamed, M.M. Elhefnawi, and N.M. Hamdy. All authors read and approved the final manuscript.

## Acknowledgements

The authors would like to acknowledge Dr. Asmaa El-Fiky, Dr. Shaimaa Yahia, and Dr. Ahmed Ragab (NRC) for their support. We also gratefully acknowledge the use of publicly available datasets that made this study possible. Specifically, we thank The Cancer Genome Atlas (TCGA) Research Network and the Genotype-Tissue Expression (GTEx) project for RNA-seq and clinical data, the International Cancer Genome Consortium (ICGC) for the LIRI-JP cohort, and the NCBI Gene Expression Omnibus (GEO) for the GSE54236 and GSE14520 microarray datasets.

## List of abbreviations

AKT: Protein Kinase B
BCL2: B-cell Lymphoma 2
BIRC5: Baculoviral Inhibitor of Apoptosis Repeat-Containing 5
CAS9: CRISPR-Associated Protein 9
CASC5: Cancer Susceptibility Candidate 5
CCA: Canonical Correlation Analysis
CCNB2: Cyclin B2
CD8: Cluster of differentiation 8
CDC20: Cell Division Cycle Protein 20 Homolog
CDK1: Cyclin Dependent Kinase 1
CHEK1: Checkpoint Kinase 1
CIBERSORT: Cell-type Identification by Estimating Relative Subsets of RNA Transcripts
CIN: Chromosomal Instability
CRISPR: Clustered Regularly Interspaced Short Palindromic Repeats
CTLA: Cytotoxic T-Lymphocyte–Associated Protein 4
CXCL10: C-X-C motif chemokine ligand 10
CXCR3: C-X-C motif chemokine receptor 3
DEGs: Differentially Expressed Genes
DMEM: Dulbecco’s Modified Eagle Medium
DPBS: Dulbecco’s Phosphate-Buffered Saline
EXO1: Exonuclease 1
FAK: Focal adhesion kinase
FBS: Fetal Bovine Serum
FN1: Fibronectin 1
Grna: Guide RNA
GTEx: Genotype-Tissue Expression
HBV: Hepatitis B Virus
HCV: Hepatitis C Virus
HLA: Human leukocyte antigen
HRP: Horseradish Peroxidase
ICI: Immune Checkpoint Inhibitor
KNL1: Kinetochore Scaffold 1
KPNA2: Karyopherin Subunit Alpha 2
MAFLD: Metabolic-Associated Fatty Liver Disease
MCC: Maximal Clique Centrality
MC3: Multi-Center Mutation Calling in Multiple Cancers
MHC: Major histocompatibility complex
MKI67: Marker of Proliferation Ki-67
MSigDB: Molecular Signatures Database
PBS: Phosphate-Buffered Saline
PCA: Principal Component Analysis
PD-1: Anti Programmed Cell Death Protein 1
PD-L1: Programmed Death-Ligand 1
PI3K: Phosphoinositide 3-Kinase
PTTG1: Pituitary Tumor Transforming Gene 1
RACGAP1: Rac GTPase Activating Protein 1
RNA-seq: RNA Sequencing
SCNAs: Somatic Copy Number Alterations
SSGSEA: Single Sample Gene Set Enrichment Analysis
STRING: Search Tool for the Retrieval of Interacting Genes/Proteins
SVs: Structural Variants
TBL: Tumor Break Load
TCGA-LIHC: The Cancer Genome Atlas Liver Hepatocellular Carcinoma
TEM: Transmission Electron Microscopy
Th2: T Helper 2 Cells
TIDE: Tumor Immune Dysfunction and Exclusion
TNF: Tumor necrosis factor
TNFRSF1A: Tumor necrosis factor receptor superfamily member 1A
TOM: Topological Overlap Matrix
TP53: Tumor Protein p53
TPX2: Targeting Protein for Xenopus Kinesin-Like Protein 2
Treg: Regulatory T Cells
UMAP: Uniform Manifold Approximation and Projection
UCSC: University of California, Santa Cruz
VEGF: Vascular Endothelial Growth Factor
WGCNA: Weighted Gene Co-expression Network Analysis

---

**Supplementary Figure S9:**
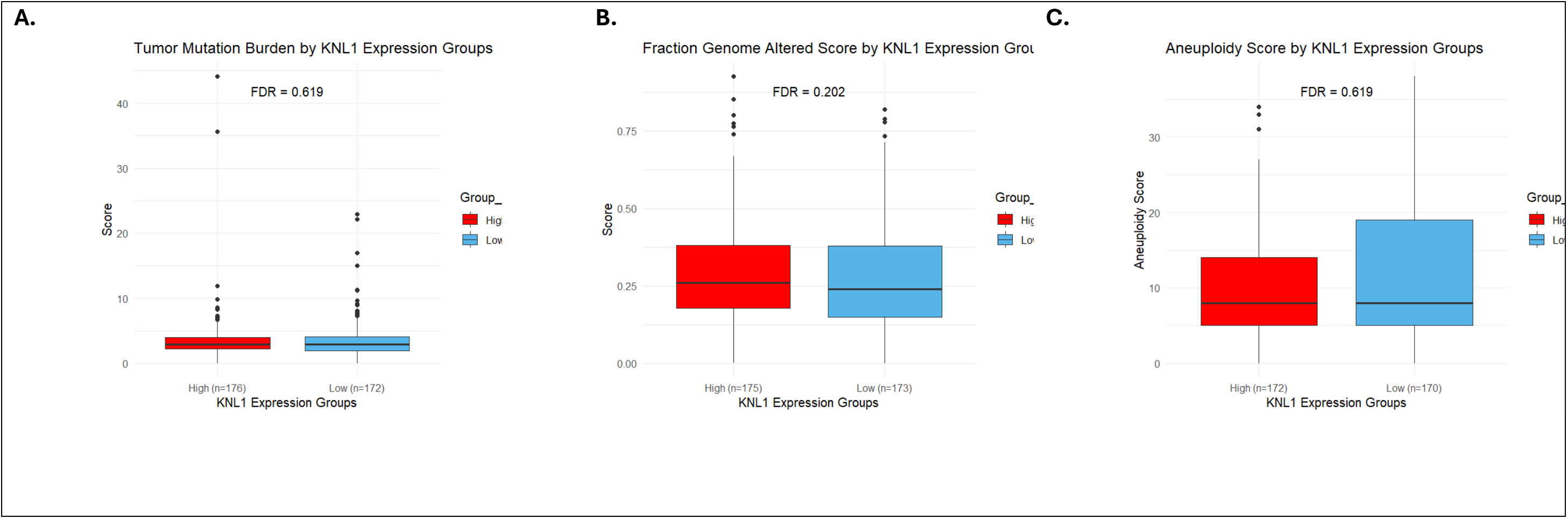
(A) Comparison of tumor mutation burden (TMB) between high and low KNL1-associated G2M network score groups (median 2.92 [IQR: 2.14–4.09] vs 2.87 [IQR: 2.03–4.00], adjusted p = 7.85 × 10⁻¹). (B) Tumor mutation burden stratified by KNL1 expression (high: median 2.90 [IQR: 2.19–4.00] vs low: median 2.90 [IQR: 1.90–4.00]). (C) Fraction of genome altered (high: median 0.26 [IQR: 0.18–0.38] vs low: median 0.23 [IQR: 0.02–0.38]). (D) Aneuploidy score (high: median 8 [IQR: 5–14] vs low: median 8 [IQR: 5–19]). Statistical comparisons were performed using the Wilcoxon rank-sum test. Data are presented as median and interquartile range (IQR). Adjusted p values were calculated using the Benjamini–Hochberg method, with adjusted p < 0.05 considered statistically significant

**Supplementary Figure S10:**
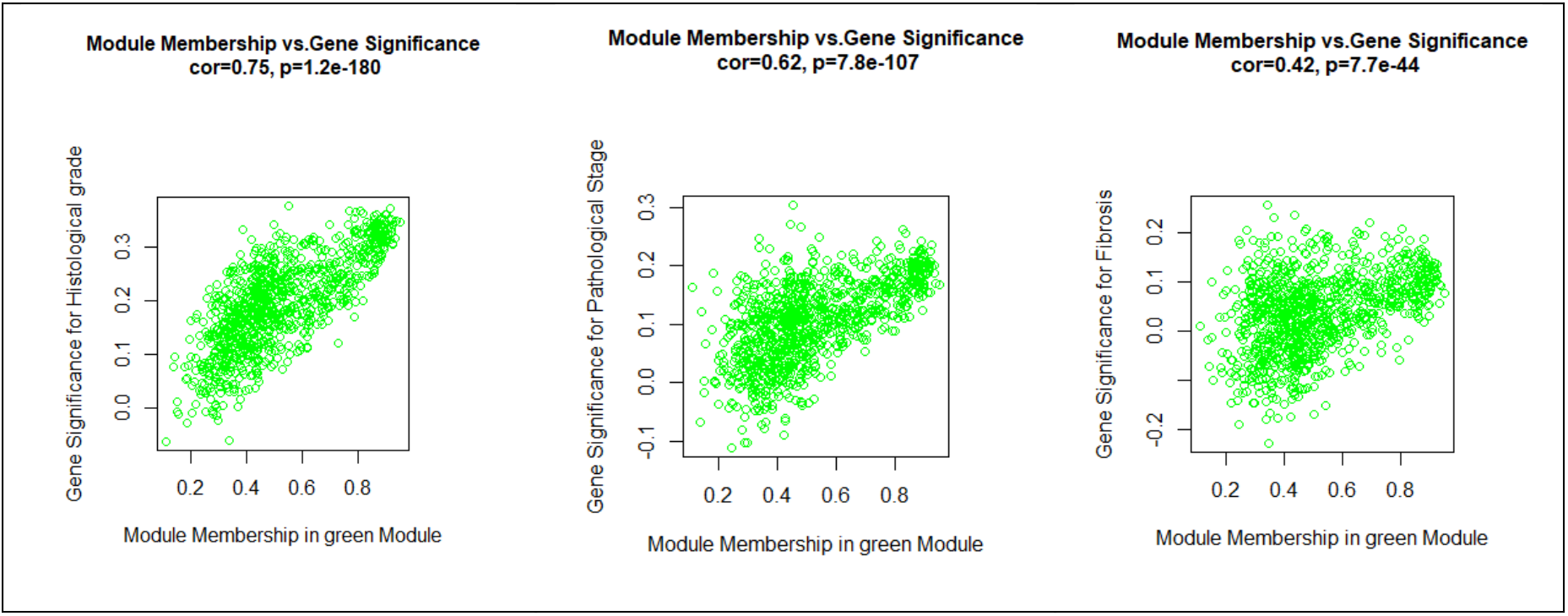
Correlation between module membership (MM) and gene significance (GS) for key clinical traits. A strong positive correlation was observed for histological grade (cor = 0.75, p = 1.2e-180), with moderate correlations for pathological stage (cor = 0.62, p = 7.8E-107) and fibrosis (cor = 0.42, p = 7.7E-44).

**Supplementary Figure S11:**
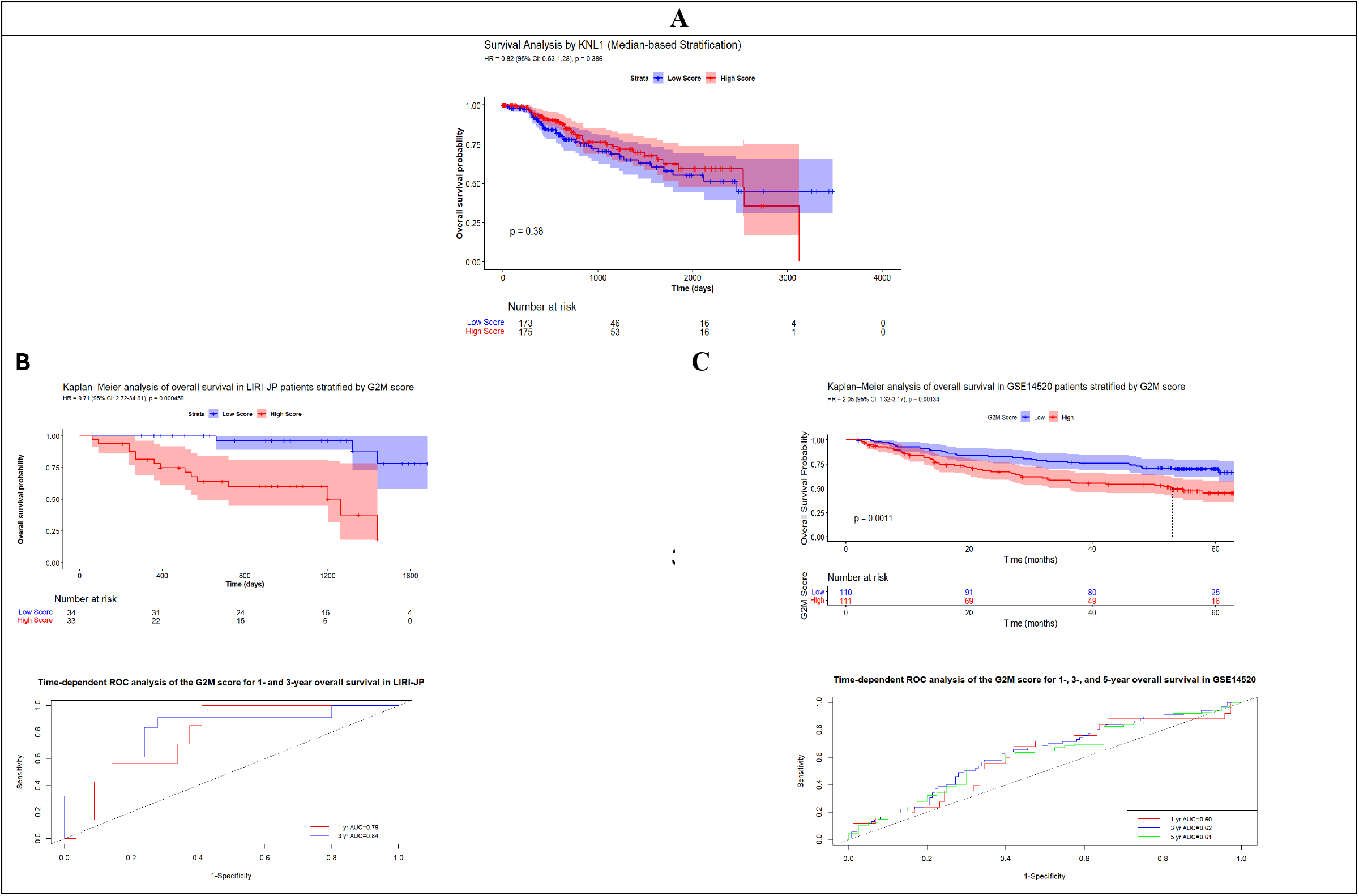
Overall Survival (OS) analyses. **(A)** Kaplan–Meier OS curves comparing patients with high (n = 175) and low (n = 173) KNL1 expression in the TCGA-LIHC cohort, dichotomized by median expression level in TCGA-LIHC cohort. **(B)** Kaplan–Meier overall survival (OS) curves comparing patients with high and low KNL1-associated G2M network scores, dichotomized by the median score in the LIRI-JP cohort (n = 67; high = 33, low = 34). Patients in the high-score group exhibited significantly poorer survival (HR = 9.71, 95% CI: 2.72–34.61, p = 4.58 × 10⁻⁴). Time-dependent ROC analysis demonstrated moderate prognostic performance of the G2M network score at 1 and 3 years (AUC = 0.79 and 0.84, respectively). **(C)** Kaplan–Meier OS curves comparing patients with high and low G2M network scores in the GSE14520 cohort (n = 225; high = 113, low = 112). Patients in the high-score group showed significantly worse survival (HR = 2.05, 95% CI: 1.32–3.17, p = 0.001). Time-dependent ROC analysis demonstrated modest predictive accuracy at 1, 3, and 5 years (AUC = 0.60, 0.62, and 0.61, respectively).

**Supplementary Figure S4:**
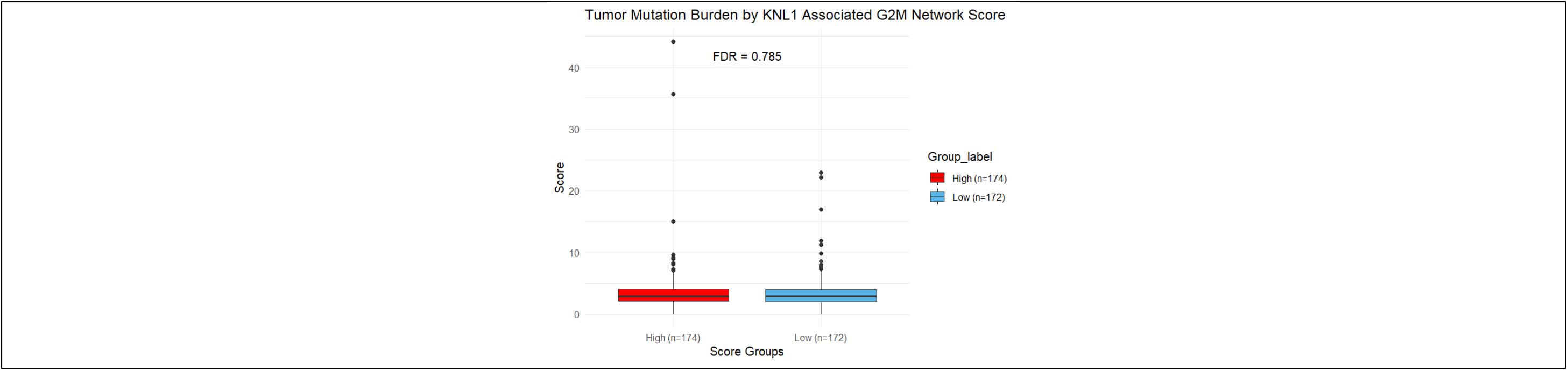
Comparison of tumor mutation burden (TMB) between high and low KNL1-associated G2M network score groups (median 2.92 [IQR: 2.14–4.09] vs 2.87 [IQR: 2.03–4.00], adjusted p = 7.85 × 10⁻¹).

## Chemiluminescence image

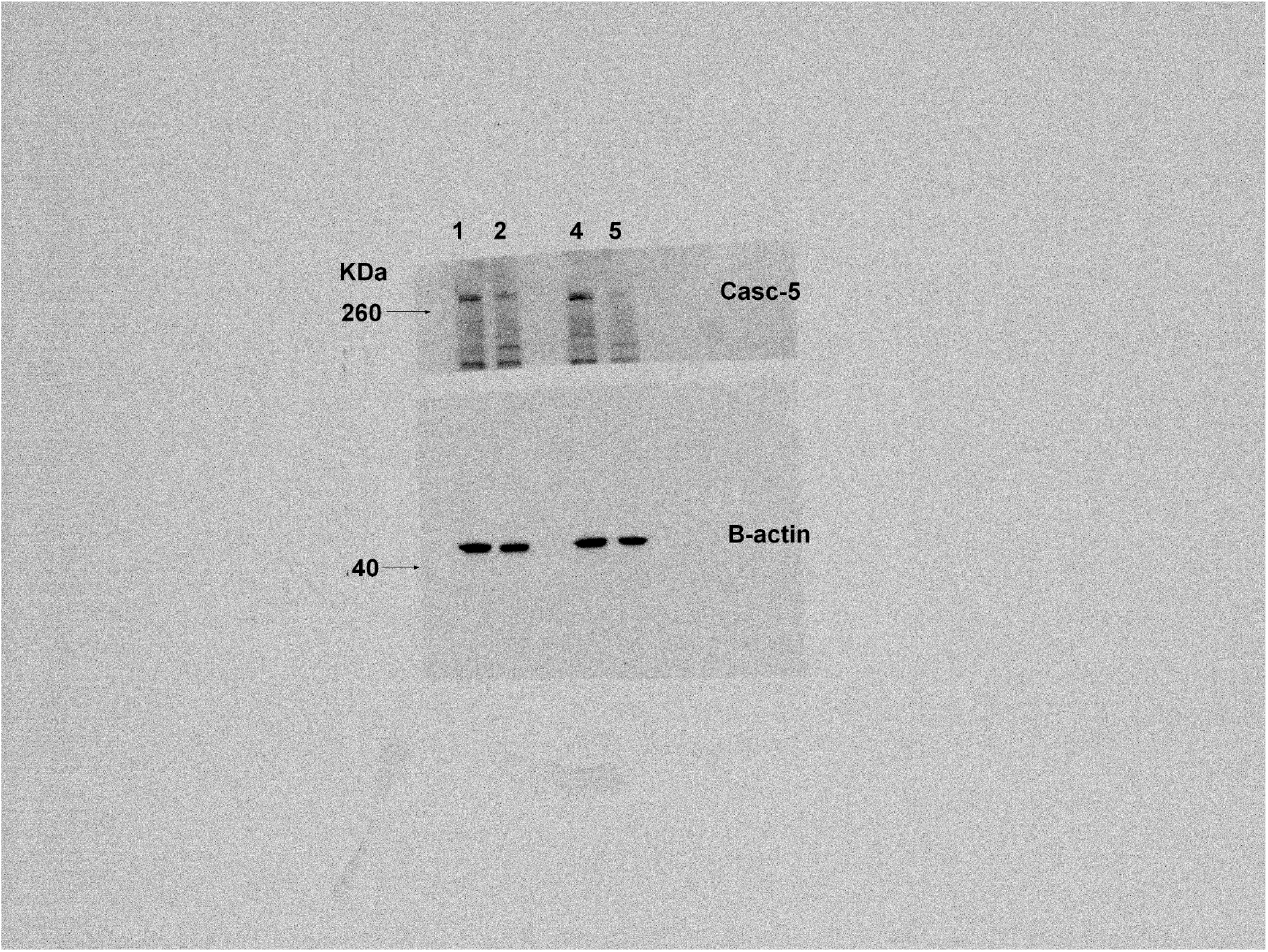

## Membrane image

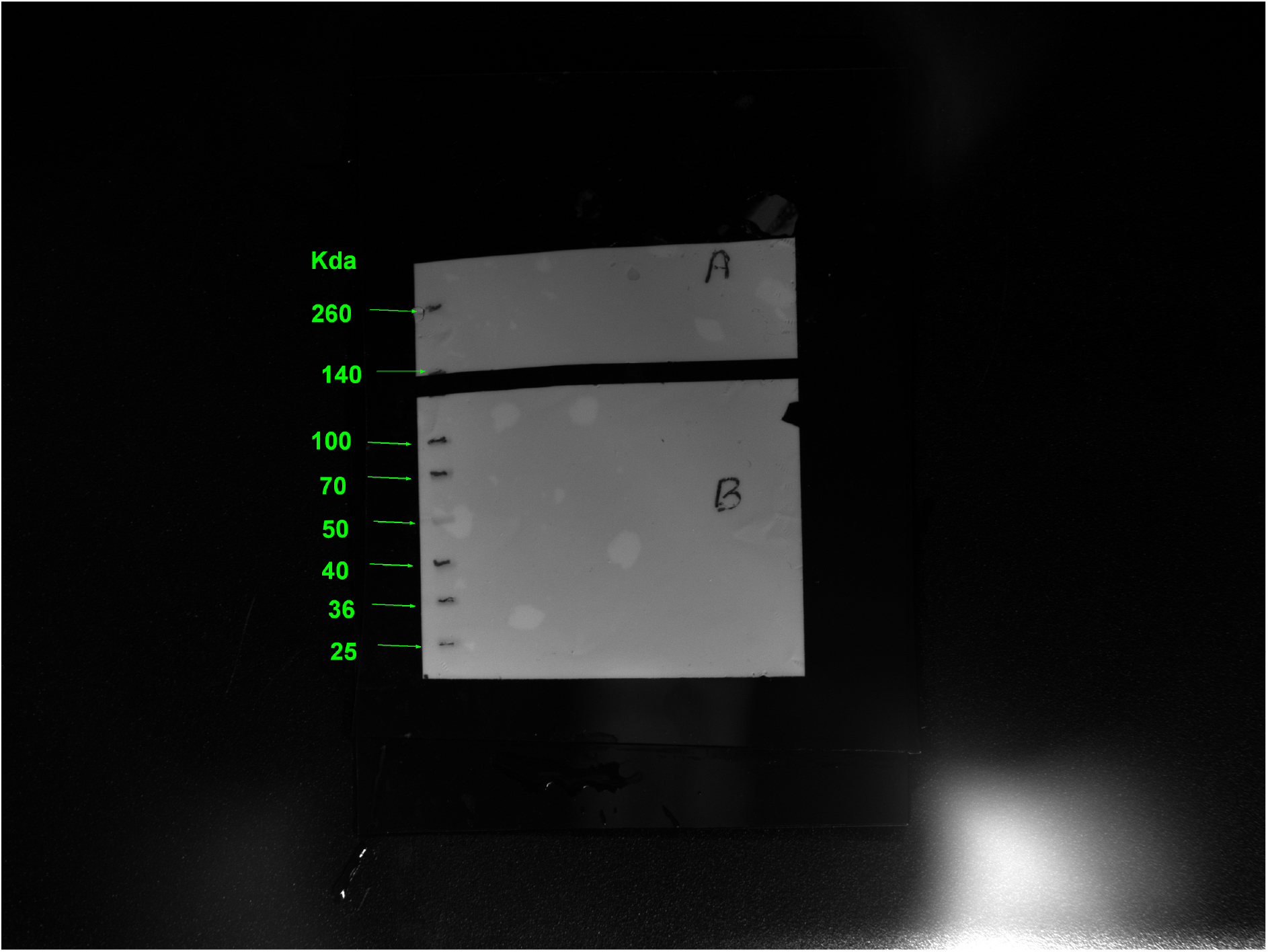

## Supplementary Tables

**Table S1:** S1-TCGA-LIHC.

| term | estimate | std.error | statistic | p.value | conf.low | conf.high |
|---|---|---|---|---|---|---|
| G2M_Score_scaled | 1.457303 | 0.148012 | 2.544299 | 0.01095 | 1.090339 | 1.947772 |
| genderMALE | 0.866762 | 0.27315 | -0.52349 | 0.600633 | 0.507451 | 1.480489 |
| age_at_initial_pathologic_diagnosis | 1.031508 | 0.011587 | 2.677236 | 0.007423 | 1.008346 | 1.055203 |
| fetoprotein_outcome_value | 0.999976 | 1.04E-05 | -2.31792 | 0.020454 | 0.999955 | 0.999996 |
| viral_status_activeHBV | 2.042591 | 0.293326 | 2.434899 | 0.014896 | 1.149481 | 3.629616 |
| viral_status_activeHCV | 2.207447 | 1.035832 | 0.764445 | 0.444602 | 0.28986 | 16.81095 |
| viral_status_activeHBV+HCV | 2.218178 | 0.473286 | 1.683308 | 0.092315 | 0.877273 | 5.60865 |
| vascular_tumor_cell_typeMicro | 1.12745 | 0.300471 | 0.399233 | 0.689722 | 0.625656 | 2.031697 |
| vascular_tumor_cell_typeMacro | 5.105766 | 0.49868 | 3.269372 | 0.001078 | 1.921249 | 13.5687 |
| neoplasm_histologic_grade | 1.456773 | 0.227334 | 1.65494 | 0.097937 | 0.933007 | 2.274567 |
| Pathologic_Stage | 1.360388 | 0.165938 | 1.854724 | 0.063636 | 0.982688 | 1.883258 |

**Table S2:** LIRI-P.

| term | estimate | std.error | statistic | p.value | conf.low | conf.high |
|---|---|---|---|---|---|---|
| G2M_scaled | 4.036632 | 0.457121 | 3.052605 | 0.002269 | 1.647847 | 9.888294 |
| donor_age_at_diagnosis.x | 1.007721 | 0.029854 | 0.25765 | 0.796677 | 0.950449 | 1.068445 |
| donor_sex.xmale | 0.346438 | 0.632577 | -1.67577 | 0.093784 | 0.100271 | 1.196949 |
| tumour_gradeI | 4.081135 | 1.28532 | 1.094183 | 0.273875 | 0.328636 | 50.68121 |
| tumour_gradeI-II | 2.612087 | 1.246125 | 0.770508 | 0.440998 | 0.227135 | 30.03939 |
| tumour_gradeII | 1.815477 | 0.835982 | 0.713351 | 0.475629 | 0.352697 | 9.345014 |
| tumour_gradeII-III | 1.27286 | 0.868967 | 0.277647 | 0.781283 | 0.2318 | 6.98951 |
| tumour_gradeIII | 5.192395 | 0.907352 | 1.815388 | 0.069464 | 0.877059 | 30.74021 |
| tumour_stage | 1.996713 | 0.318028 | 2.174346 | 0.029679 | 1.070557 | 3.7241 |

**Table S3:** GSE4520.

| term | estimate | std.error | statistic | p.value | conf.low | conf.high |
|---|---|---|---|---|---|---|
| G2M_Score_scaled | 1.348401 | 0.125932 | 2.373661 | 0.017613 | 1.053479 | 1.725887 |
| Age | 0.996169 | 0.010118 | -0.37934 | 0.704437 | 0.976609 | 1.016121 |
| GenderM | 1.729582 | 0.373222 | 1.467974 | 0.142111 | 0.832251 | 3.594415 |
| AFPhigh | 1.476662 | 0.225006 | 1.732327 | 0.083215 | 0.95007 | 2.295128 |
| HBV_simpleHBV_positive | 1.123445 | 0.718918 | 0.16191 | 0.871377 | 0.274541 | 4.597222 |

## References

1. International Agency for Research on Cancer TIA for R on. Global Cancer Observatory [Internet]. 2022 [cited 2025 Aug 15]. GLOBOCAN 2022: Liver Cancer Fact Sheet. Available from: https://gco.iarc.fr/

2. Bray F, Laversanne M, Sung H, Ferlay J, Siegel RL, Soerjomataram I, et al. Global cancer statistics 2022: GLOBOCAN estimates of incidence and mortality worldwide for 36 cancers in 185 countries. CA Cancer J Clin. 2024;74(3):229–63. doi:10.3322/caac.21834

3. Llovet JM, Kelley RK, Villanueva A, Singal AG, Pikarsky E, Roayaie S, et al. Hepatocellular carcinoma. Nat Rev Dis Primer. 2021 Jan 21;7(1):1–28. doi:10.1038/s41572-020-00240-3

4. Swellam M, Hamdy NM. Association of nonalcoholic fatty liver disease with a single nucleotide polymorphism on the gene encoding leptin receptor. IUBMB Life. 2012 Feb;64(2):180–6. doi:10.1002/iub.597

5. Reig M, Forner A, Rimola J, Ferrer-Fàbrega J, Burrel M, Garcia-Criado Á, et al. BCLC strategy for prognosis prediction and treatment recommendation: The 2022 update. J Hepatol. 2022 Mar 1;76(3):681–93. doi:10.1016/j.jhep.2021.11.018 PubMed PMID: 34801630.

6. Abou-Alfa GK, Meyer T, Cheng AL, El-Khoueiry AB, Rimassa L, Ryoo BY, et al. Cabozantinib in Patients with Advanced and Progressing Hepatocellular Carcinoma. N Engl J Med. 2018 Jul 5;379(1):54–63. doi:10.1056/NEJMoa1717002 PubMed PMID: 29972759; PubMed Central PMCID: PMC7523244.

7. Bruix J, Qin S, Merle P, Granito A, Huang YH, Bodoky G, et al. Regorafenib for patients with hepatocellular carcinoma who progressed on sorafenib treatment (RESORCE): a randomised, double-blind, placebo-controlled, phase 3 trial. Lancet Lond Engl. 2017 Jan 7;389(10064):56–66. doi:10.1016/S0140-6736(16)32453-9 PubMed PMID: 27932229.

8. Kudo M, Finn RS, Qin S, Han KH, Ikeda K, Piscaglia F, et al. Lenvatinib versus sorafenib in first-line treatment of patients with unresectable hepatocellular carcinoma: a randomised phase 3 non-inferiority trial. Lancet Lond Engl. 2018 Mar 24;391(10126):1163–73. doi:10.1016/S0140-6736(18)30207-1 PubMed PMID: 29433850.

9. Ladd AD, Duarte S, Sahin I, Zarrinpar A. Mechanisms of drug resistance in HCC. Hepatology. 2024 Apr;79(4):926. doi:10.1097/HEP.0000000000000237

10. Llovet JM, Ricci S, Mazzaferro V, Hilgard P, Gane E, Blanc JF, et al. Sorafenib in advanced hepatocellular carcinoma. N Engl J Med. 2008 Jul 24;359(4):378–90. doi:10.1056/NEJMoa0708857 PubMed PMID: 18650514.

11. Zhu AX, Kang YK, Yen CJ, Finn RS, Galle PR, Llovet JM, et al. Ramucirumab after sorafenib in patients with advanced hepatocellular carcinoma and increased α-fetoprotein concentrations (REACH-2): a randomised, double-blind, placebo-controlled, phase 3 trial. Lancet Oncol. 2019 Feb;20(2):282–96. doi:10.1016/S1470-2045(18)30937-9 PubMed PMID: 30665869.

12. Finn RS, Qin S, Ikeda M, Galle PR, Ducreux M, Kim TY, et al. Atezolizumab plus Bevacizumab in Unresectable Hepatocellular Carcinoma. N Engl J Med. 2020 May 14;382(20):1894–905. doi:10.1056/NEJMoa1915745 PubMed PMID: 32402160.

13. Abou-Alfa GK, Lau G, Kudo M, Chan SL, Kelley RK, Furuse J, et al. Tremelimumab plus Durvalumab in Unresectable Hepatocellular Carcinoma. NEJM Evid. 2022 Jul 26;1(8). doi:10.1056/EVIDoa2100070

14. Yau T, Park JW, Finn RS, Cheng AL, Mathurin P, Edeline J, et al. Nivolumab versus sorafenib in advanced hepatocellular carcinoma (CheckMate 459): a randomised, multicentre, open-label, phase 3 trial. Lancet Oncol. 2022 Jan;23(1):77–90. doi:10.1016/S1470-2045(21)00604-5 PubMed PMID: 34914889.

15. Abdelhamed W, El-Kassas M. Hepatocellular carcinoma recurrence: Predictors and management. Liver Res. 2023 Dec;7(4):321–32. doi:10.1016/j.livres.2023.11.004

16. Hamdy NM, Basalious EB, El-Sisi MG, Nasr M, Kabel AM, Nossier ES, et al. Advancements in current one-size-fits-all therapies compared to future treatment innovations for better improved chemotherapeutic outcomes: a step-toward personalized medicine. Curr Med Res Opin. 2024 Nov;40(11):1943–61. doi:10.1080/03007995.2024.2416985 PubMed PMID: 39412377.

17. Gomaa A, Waked I. Management of advanced hepatocellular carcinoma: review of current and potential therapies. Hepatoma Res. 2017 Jun 15;3(0):112–22. doi:10.20517/2394-5079.2017.03

18. Park JW, Chen M, Colombo M, Roberts LR, Schwartz M, Chen PJ, et al. Global patterns of hepatocellular carcinoma management from diagnosis to death: the BRIDGE Study. Liver Int Off J Int Assoc Study Liver. 2015 Sep;35(9):2155–66. doi:10.1111/liv.12818 PubMed PMID: 25752327; PubMed Central PMCID: PMC4691343.

19. Safri F, Nguyen R, Zerehpooshnesfchi S, George J, Qiao L. Heterogeneity of hepatocellular carcinoma: from mechanisms to clinical implications. Cancer Gene Ther. 2024;31(8):1105–12. doi:10.1038/s41417-024-00764-w PubMed PMID: 38499648; PubMed Central PMCID: PMC11327108.

20. Khalil BM, Shahin MH, Solayman MHM, Langaee T, Schaalan MF, Gong Y, et al. Genetic and Nongenetic Factors Affecting Clopidogrel Response in the Egyptian Population. Clin Transl Sci. 2016 Feb;9(1):23–8. doi:10.1111/cts.12383 PubMed PMID: 26757134; PubMed Central PMCID: PMC4760893.

21. Prakash J, Shaked Y. The Interplay between Extracellular Matrix Remodeling and Cancer Therapeutics. Cancer Discov. 2024 Aug 2;14(8):1375–88. doi:10.1158/2159-8290.CD-24-0002 PubMed PMID: 39091205; PubMed Central PMCID: PMC11294818.

22. Hu Q, Zhu Y, Mei J, Liu Y, Zhou G. Extracellular matrix dynamics in tumor immunoregulation: from tumor microenvironment to immunotherapy. J Hematol OncolJ Hematol Oncol. 2025 Jun 19;18:65. doi:10.1186/s13045-025-01717-y PubMed PMID: 40537775; PubMed Central PMCID: PMC12178005.

23. Yuan Z, Li Y, Zhang S, Wang X, Dou H, Yu X, et al. Extracellular matrix remodeling in tumor progression and immune escape: from mechanisms to treatments. Mol Cancer. 2023 Mar 11;22:48. doi:10.1186/s12943-023-01744-8 PubMed PMID: 36906534; PubMed Central PMCID: PMC10007858.

24. Hamdy NM, Sallam AAM, Elazazy O, Kabel AM, Salama RM, Gouhar SA, et al. LincRNA-miR interactions in hepatocellular carcinoma: comprehensive review and in silico analysis: a step toward ncRNA precision. Naunyn Schmiedebergs Arch Pharmacol. 2025 May 23. doi:10.1007/s00210-025-04285-7 PubMed PMID: 40410550.

25. Youness RA, Hassan HA, Abaza T, Hady AA, El Magdoub HM, Ali M, et al. A Comprehensive Insight and In Silico Analysis of CircRNAs in Hepatocellular Carcinoma: A Step toward ncRNA-Based Precision Medicine. Cells. 2024 Jul 24;13(15):1245. doi:10.3390/cells13151245 PubMed PMID: 39120276; PubMed Central PMCID: PMC11312109.

26. Chan LK, Tsui YM, Ho DWH, Ng IOL. Cellular heterogeneity and plasticity in liver cancer. Semin Cancer Biol. 2022 Jul;82:134–49. doi:10.1016/j.semcancer.2021.02.015 PubMed PMID: 33647386.

27. Atta H, Alzahaby N, Hamdy NM, Emam SH, Sonousi A, Ziko L. New trends in synthetic drugs and natural products targeting 20S proteasomes in cancers. Bioorganic Chem. 2023 Apr;133:106427. doi:10.1016/j.bioorg.2023.106427 PubMed PMID: 36841046.

28. Schulze K, Imbeaud S, Letouzé E, Alexandrov LB, Calderaro J, Rebouissou S, et al. Exome sequencing of hepatocellular carcinomas identifies new mutational signatures and potential therapeutic targets. Nat Genet. 2015 May;47(5):505–11. doi:10.1038/ng.3252

29. Yang C, Zhang S, Cheng Z, Liu Z, Zhang L, Jiang K, et al. Multi-region sequencing with spatial information enables accurate heterogeneity estimation and risk stratification in liver cancer. Genome Med. 2022 Dec 16;14(1):142. doi:10.1186/s13073-022-01143-6 PubMed PMID: 36527145; PubMed Central PMCID: PMC9758830.

30. Bach DH, Zhang W, Sood AK. Chromosomal Instability in Tumor Initiation and Development. Cancer Res. 2019 Aug 15;79(16):3995–4002. doi:10.1158/0008-5472.CAN-18-3235 PubMed PMID: 31350294; PubMed Central PMCID: PMC7694409.

31. Ebert P, Audano PA, Zhu Q, Rodriguez-Martin B, Porubsky D, Bonder MJ, et al. Haplotype-resolved diverse human genomes and integrated analysis of structural variation. Science. 2021 Apr 2;372(6537):eabf7117. doi:10.1126/science.abf7117 PubMed PMID: 33632895; PubMed Central PMCID: PMC8026704.

32. Hamdan A, Ewing A. Unravelling the tumour genome: The evolutionary and clinical impacts of structural variants in tumourigenesis. J Pathol. 2022 Jul;257(4):479–93. doi:10.1002/path.5901 PubMed PMID: 35355264; PubMed Central PMCID: PMC9321913.

33. Mahmoud M, Gobet N, Cruz-Dávalos DI, Mounier N, Dessimoz C, Sedlazeck FJ. Structural variant calling: the long and the short of it. Genome Biol. 2019 Nov 20;20(1):246. doi:10.1186/s13059-019-1828-7 PubMed PMID: 31747936; PubMed Central PMCID: PMC6868818.

34. Yarchoan M, Hopkins A, Jaffee EM. Tumor Mutational Burden and Response Rate to PD-1 Inhibition. N Engl J Med. 2017 Dec 21;377(25):2500–1. doi:10.1056/NEJMc1713444 PubMed PMID: 29262275; PubMed Central PMCID: PMC6549688.

35. Samstein RM, Lee CH, Shoushtari AN, Hellmann MD, Shen R, Janjigian YY, et al. Tumor mutational load predicts survival after immunotherapy across multiple cancer types. Nat Genet. 2019 Feb;51(2):202–6. doi:10.1038/s41588-018-0312-8 PubMed PMID: 30643254; PubMed Central PMCID: PMC6365097.

36. Vasudevan A, Schukken KM, Sausville EL, Girish V, Adebambo OA, Sheltzer JM. Aneuploidy as a promoter and suppressor of malignant growth. Nat Rev Cancer. 2021 Feb;21(2):89–103. doi:10.1038/s41568-020-00321-1 PubMed PMID: 33432169.

37. Lakbir S, Lahoz S, Cuatrecasas M, Camps J, Glas RA, Heringa J, et al. Tumour break load is a biologically relevant feature of genomic instability with prognostic value in colorectal cancer. Eur J Cancer. 2022 Dec;177:94–102. doi:10.1016/j.ejca.2022.09.034 PubMed PMID: 36334560.

38. Yuen KW, Montpetit B, Hieter P. The kinetochore and cancer: what’s the connection? Curr Opin Cell Biol. 2005 Dec;17(6):576–82. doi:10.1016/j.ceb.2005.09.012

39. Caldas GV, DeLuca JG. KNL1: Bringing Order to the Kinetochore. Chromosoma. 2014 Jun;123(3):169–81. doi:10.1007/s00412-013-0446-5 PubMed PMID: 24310619; PubMed Central PMCID: PMC4032621.

40. Ding Y, Wang K, Zhao S, Li Y, Qiu W, Zhu C, et al. Role of Kinetochore Scaffold 1 (KNL1) in Tumorigenesis and Tumor Immune Microenvironment in Pan-Cancer: Bioinformatics Analyses and Validation of Expression. Int J Gen Med. 2023 Oct 31;16:4883–906. doi:10.2147/IJGM.S424245 PubMed PMID: 37928953; PubMed Central PMCID: PMC10625436.

41. Zhang Y, Ji Q, Wang J, Dong Y, Pang M, Fu S, et al. High expression of KNL1 in prostate adenocarcinoma is associated with poor prognosis and immune infiltration. Front Genet. 2023 Jan 6;13. doi:10.3389/fgene.2022.1100787

42. Dong Y, Wu T, Chen J, Mo L, You Y. KNL1 is a lung cancer prognostic biomarker associated with the immune microenvironment [Internet]. Research Square; 2024 [cited 2025 May 16]. Available from: https://www.researchsquare.com/article/rs-4379762/v1 doi:10.21203/rs.3.rs-4379762/v1

43. Cui Y, Zhang C, Ma S, Guo W, Cao W, Guan F. CASC5 is a potential tumour driving gene in lung adenocarcinoma. Cell Biochem Funct. 2020 Aug;38(6):733–42. doi:10.1002/cbf.3540 PubMed PMID: 32283571.

44. Singh PK, Bhatt MLB, Singh P, Rath SK, Dalela D, Goel MM. CASC5 is a potential cancer-testis gene in human urinary bladder transitional cell carcinoma. Drug Discov Ther. 2021;15(6):331–6. doi:10.5582/ddt.2021.01108 PubMed PMID: 35034926.

45. Song B, Du J, Song DF, Ren JC, Feng Y. Dysregulation of NCAPG, KNL1, miR-148a-3p, miR-193b-3p, and miR-1179 may contribute to the progression of gastric cancer. Biol Res. 2018 Nov 3;51(1):44. doi:10.1186/s40659-018-0192-5 PubMed PMID: 30390708; PubMed Central PMCID: PMC6215350.

46. He K, Li J, Huang X, Zhao W, Wang K, Wang T, et al. KNL1 is a prognostic and diagnostic biomarker related to immune infiltration in patients with uterine corpus endometrial carcinoma. Front Oncol. 2023 Jan 27;13:1090779. doi:10.3389/fonc.2023.1090779 PubMed PMID: 36776306; PubMed Central PMCID: PMC9913269.

47. Nevi L, Aiello C, Molinaro F, Altomare A, Musarra T, Gallo P, et al. Decoding the molecular and genomic landscape of hepatocellular carcinoma: biomarker discovery, classification frameworks, and therapeutic targeting. Npj Gut Liver. 2025 Oct 1;2(1):25. doi:10.1038/s44355-025-00038-8

48. Wang S, Liu X. The UCSCXenaTools R package: a toolkit for accessing genomics data from UCSC Xena platform, from cancer multi-omics to single-cell RNA-seq. J Open Source Softw. 2019 Aug 5;4(40):1627. doi:10.21105/joss.01627

49. Vivian J, Rao AA, Nothaft FA, Ketchum C, Armstrong J, Novak A, et al. Toil enables reproducible, open source, big biomedical data analyses. Nat Biotechnol. 2017 Apr 11;35(4):314–6. doi:10.1038/nbt.3772 PubMed PMID: 28398314; PubMed Central PMCID: PMC5546205.

50. Cerami E, Gao J, Dogrusoz U, Gross BE, Sumer SO, Aksoy BA, et al. The cBio cancer genomics portal: an open platform for exploring multidimensional cancer genomics data. Cancer Discov. 2012 May;2(5):401–4. doi:10.1158/2159-8290.CD-12-0095 PubMed PMID: 22588877; PubMed Central PMCID: PMC3956037.

51. Chen HM, MacDonald JA. Network analysis of TCGA and GTEx gene expression datasets for identification of trait-associated biomarkers in human cancer. STAR Protoc. 2022 Mar;3(1):101168. doi:10.1016/j.xpro.2022.101168

52. Chen Y, Chen L, Lun ATL, Baldoni PL, Smyth GK. edgeR v4: powerful differential analysis of sequencing data with expanded functionality and improved support for small counts and larger datasets. Nucleic Acids Res. 2025 Jan 11;53(2):gkaf018. doi:10.1093/nar/gkaf018 PubMed PMID: 39844453; PubMed Central PMCID: PMC11754124.

53. Law CW, Chen Y, Shi W, Smyth GK. voom: precision weights unlock linear model analysis tools for RNA-seq read counts. Genome Biol. 2014 Feb 3;15(2):R29. doi:10.1186/gb-2014-15-2-r29

54. Ritchie ME, Phipson B, Wu D, Hu Y, Law CW, Shi W, et al. limma powers differential expression analyses for RNA-sequencing and microarray studies. Nucleic Acids Res. 2015 Jan 20;43(7):e47. doi:10.1093/nar/gkv007 PubMed PMID: 25605792.

55. Gao J, Aksoy BA, Dogrusoz U, Dresdner G, Gross B, Sumer SO, et al. Integrative analysis of complex cancer genomics and clinical profiles using the cBioPortal. Sci Signal. 2013 Apr 2;6(269):pl1. doi:10.1126/scisignal.2004088 PubMed PMID: 23550210; PubMed Central PMCID: PMC4160307.

56. de Bruijn I, Kundra R, Mastrogiacomo B, Tran TN, Sikina L, Mazor T, et al. Analysis and Visualization of Longitudinal Genomic and Clinical Data from the AACR Project GENIE Biopharma Collaborative in cBioPortal. Cancer Res. 2023 Dec 1;83(23):3861–7. doi:10.1158/0008-5472.CAN-23-0816 PubMed PMID: 37668528; PubMed Central PMCID: PMC10690089.

57. Donehower LA, Soussi T, Korkut A, Liu Y, Schultz A, Cardenas M, et al. Integrated Analysis of TP53 Gene and Pathway Alterations in The Cancer Genome Atlas. Cell Rep. 2019 Jul 30;28(5):1370–1384.e5. doi:10.1016/j.celrep.2019.07.001 PubMed PMID: 31365877; PubMed Central PMCID: PMC7546539.

58. Langfelder P, Horvath S. WGCNA: an R package for weighted correlation network analysis. BMC Bioinformatics. 2008 Dec 29;9:559. doi:10.1186/1471-2105-9-559 PubMed PMID: 19114008; PubMed Central PMCID: PMC2631488.

59. Langfelder P, Horvath S. Fast R Functions for Robust Correlations and Hierarchical Clustering. J Stat Softw. 2012 Mar;46(11):i11. PubMed PMID: 23050260; PubMed Central PMCID: PMC3465711.

60. Carlson M. org.Hs.eg.db: Genome wide annotation for Human [Internet]. 2024 [cited 2025 May 10]. Available from: http://bioconductor.org/packages/org.Hs.eg.db/

61. Yu G, Wang LG, Han Y, He QY. clusterProfiler: an R package for comparing biological themes among gene clusters. Omics J Integr Biol. 2012 May;16(5):284–7. doi:10.1089/omi.2011.0118 PubMed PMID: 22455463; PubMed Central PMCID: PMC3339379.

62. Xu S, Hu E, Cai Y, Xie Z, Luo X, Zhan L, et al. Using clusterProfiler to characterize multiomics data. Nat Protoc. 2024 Nov;19(11):3292–320. doi:10.1038/s41596-024-01020-z PubMed PMID: 39019974.

63. Yu G. Thirteen years of clusterProfiler. The Innovation. 2024 Nov 4;5(6):100722. doi:10.1016/j.xinn.2024.100722

64. Dolgalev I. msigdbr: MSigDB Gene Sets for Multiple Organisms in a Tidy Data Format [Internet]. 2018 [cited 2025 May 10]. p. 10.0.2. Available from: https://CRAN.R-project.org/package=msigdbr doi:10.32614/CRAN.package.msigdbr

65. Igor Dolgalev. msigdbr: MSigDB Gene Sets for Multiple Organisms in a Tidy Data Format [R package version 25.1.1]. 2025. Available from: https://github.com/igordot/msigdbr

66. Shannon P, Markiel A, Ozier O, Baliga NS, Wang JT, Ramage D, et al. Cytoscape: a software environment for integrated models of biomolecular interaction networks. Genome Res. 2003 Nov;13(11):2498–504. doi:10.1101/gr.1239303 PubMed PMID: 14597658; PubMed Central PMCID: PMC403769.

67. Doncheva NT, Morris JH, Gorodkin J, Jensen LJ. Cytoscape StringApp: Network Analysis and Visualization of Proteomics Data. J Proteome Res. 2019;18(2):623–32. doi:10.1021/acs.jproteome.8b00702

68. Linlin Yan. ggvenn: Draw Venn Diagram by ’ggplot2 [Internet]. 2023. Available from: https://CRAN.R-project.org/package=ggvenn

69. Szklarczyk D, Franceschini A, Wyder S, Forslund K, Heller D, Huerta-Cepas J, et al. STRING v10: protein-protein interaction networks, integrated over the tree of life. Nucleic Acids Res. 2015 Jan;43(Database issue):D447–452. doi:10.1093/nar/gku1003 PubMed PMID: 25352553; PubMed Central PMCID: PMC4383874.

70. Chin CH, Chen SH, Wu HH, Ho CW, Ko MT, Lin CY. cytoHubba: Identifying hub objects and sub-networks from complex interactome. BMC Syst Biol. 2014;8(4):1–7. doi:10.1186/1752-0509-8-S4-S11

71. Hänzelmann S, Castelo R, Guinney J. GSVA: gene set variation analysis for microarray and RNA-seq data. BMC Bioinformatics. 2013 Jan 16;14:7. doi:10.1186/1471-2105-14-7 PubMed PMID: 23323831; PubMed Central PMCID: PMC3618321.

72. Gu Z, Eils R, Schlesner M. Complex heatmaps reveal patterns and correlations in multidimensional genomic data. Bioinformatics. 2016 Sep 15;32(18):2847–9. doi:10.1093/bioinformatics/btw313

73. Alboukadel Kassambara, Marcin Kosinski, Przemyslaw Biecek. survminer: Drawing Survival Curves using “ggplot2” [Internet]. 2024. Available from: https://CRAN.R-project.org/package=survminer

74. Therneau TM, Grambsch PM. Modeling Survival Data: Extending the Cox Model [Internet]. New York, NY: Springer; 2000 [cited 2025 Dec 16]. (Dietz K, Gail M, Krickeberg K, Samet J, Tsiatis A, editors. Statistics for Biology and Health). Available from: http://link.springer.com/10.1007/978-1-4757-3294-8 doi:10.1007/978-1-4757-3294-8

75. Blanche P, Dartigues JF, Jacqmin-Gadda H. Estimating and comparing time-dependent areas under receiver operating characteristic curves for censored event times with competing risks. Stat Med. 2013 Dec 30;32(30):5381–97. doi:10.1002/sim.5958 PubMed PMID: 24027076.

76. Zhang J, Bajari R, Andric D, Gerthoffert F, Lepsa A, Nahal-Bose H, et al. The International Cancer Genome Consortium Data Portal. Nat Biotechnol. 2019 Apr;37(4):367–9. doi:10.1038/s41587-019-0055-9 PubMed PMID: 30877282.

77. Villa E, Critelli R, Lei B, Marzocchi G, Cammà C, Giannelli G, et al. Neoangiogenesis-related genes are hallmarks of fast-growing hepatocellular carcinomas and worst survival. Results from a prospective study. Gut. 2016 May;65(5):861–9. doi:10.1136/gutjnl-2014-308483 PubMed PMID: 25666192.

78. Roessler S, Jia HL, Budhu A, Forgues M, Ye QH, Lee JS, et al. A unique metastasis gene signature enables prediction of tumor relapse in early-stage hepatocellular carcinoma patients. Cancer Res. 2010 Dec 15;70(24):10202–12. doi:10.1158/0008-5472.CAN-10-2607 PubMed PMID: 21159642; PubMed Central PMCID: PMC3064515.

79. Davis S, Meltzer PS. GEOquery: a bridge between the Gene Expression Omnibus (GEO) and BioConductor. Bioinforma Oxf Engl. 2007 Jul 15;23(14):1846–7. doi:10.1093/bioinformatics/btm254 PubMed PMID: 17496320.

80. Wickham H. ggplot2 [Internet]. Cham: Springer International Publishing; 2016 [cited 2025 Aug 1]. (Use R!). Available from: http://link.springer.com/10.1007/978-3-319-24277-4 doi:10.1007/978-3-319-24277-4

81. Thorsson V, Gibbs DL, Brown SD, Wolf D, Bortone DS, Yang THO, et al. The Immune Landscape of Cancer. Immunity. 2018 Apr 17;48(4):812–830.e14. doi:10.1016/j.immuni.2018.03.023 PubMed PMID: 29628290.

82. Newman AM, Liu CL, Green MR, Gentles AJ, Feng W, Xu Y, et al. Robust enumeration of cell subsets from tissue expression profiles. Nat Methods. 2015 May;12(5):453–7. doi:10.1038/nmeth.3337

83. Jiang P, Gu S, Pan D, Fu J, Sahu A, Hu X, et al. Signatures of T cell dysfunction and exclusion predict cancer immunotherapy response. Nat Med. 2018 Oct;24(10):1550–8. doi:10.1038/s41591-018-0136-1 PubMed PMID: 30127393; PubMed Central PMCID: PMC6487502.

84. Rhee JK, Jung YC, Kim KR, Yoo J, Kim J, Lee YJ, et al. Impact of Tumor Purity on Immune Gene Expression and Clustering Analyses across Multiple Cancer Types. Cancer Immunol Res. 2018 Jan;6(1):87–97. doi:10.1158/2326-6066.CIR-17-0201 PubMed PMID: 29141981.

85. Ahn S, Grimes T, Datta S. The Analysis of Gene Expression Data Incorporating Tumor Purity Information. Front Genet. 2021 Aug 23;12:642759. doi:10.3389/fgene.2021.642759 PubMed PMID: 34497631; PubMed Central PMCID: PMC8419469.

86. Sasiain I, Nacer DF, Aine M, Veerla S, Staaf J. Tumor purity estimated from bulk DNA methylation can be used for adjusting beta values of individual samples to better reflect tumor biology. NAR Genomics Bioinforma. 2024 Nov 4;6(4):lqae146. doi:10.1093/nargab/lqae146 PubMed PMID: 39498434; PubMed Central PMCID: PMC11532792.

87. Butler A, Hoffman P, Smibert P, Papalexi E, Satija R. Integrating single-cell transcriptomic data across different conditions, technologies, and species. Nat Biotechnol. 2018 May;36(5):411–20. doi:10.1038/nbt.4096

88. Stuart T, Butler A, Hoffman P, Hafemeister C, Papalexi E, Mauck WM, et al. Comprehensive Integration of Single-Cell Data. Cell. 2019 Jun 13;177(7):1888–1902.e21. doi:10.1016/j.cell.2019.05.031 PubMed PMID: 31178118; PubMed Central PMCID: PMC6687398.

89. Hao Y, Hao S, Andersen-Nissen E, Mauck W, Zheng S, Butler A, et al. Integrated analysis of multimodal single-cell dat [Internet]. 2020 [cited 2025 Aug 1]. Available from: https://europepmc.org/article/PPR/PPR224971 doi:10.1101/2020.10.12.335331

90. Hao Y, Stuart T, Kowalski MH, Choudhary S, Hoffman P, Hartman A, et al. Dictionary learning for integrative, multimodal and scalable single-cell analysis. Nat Biotechnol. 2024 Feb;42(2):293–304. doi:10.1038/s41587-023-01767-y PubMed PMID: 37231261; PubMed Central PMCID: PMC10928517.

91. Lu Y, Yang A, Quan C, Pan Y, Zhang H, Li Y, et al. A single-cell atlas of the multicellular ecosystem of primary and metastatic hepatocellular carcinoma. Nat Commun. 2022 Aug 6;13(1):4594. doi:10.1038/s41467-022-32283-3

92. Xu J, Cui L, Zhuang J, Meng Y, Bing P, He B, et al. Evaluating the performance of dropout imputation and clustering methods for single-cell RNA sequencing data. Comput Biol Med. 2022 Jul;146:105697. doi:10.1016/j.compbiomed.2022.105697

93. Jin S, Guerrero-Juarez CF, Zhang L, Chang I, Ramos R, Kuan CH, et al. Inference and analysis of cell-cell communication using CellChat. Nat Commun. 2021 Feb 17;12(1):1088. doi:10.1038/s41467-021-21246-9

94. Suoqin Jin. Inference and analysis of cell-cell communication using CellChat [R package version 1.6.1]. 2025. Available from: https://github.com/sqjin/CellChat

95. Javaid N, Choi S. CRISPR/Cas System and Factors Affecting Its Precision and Efficiency. Front Cell Dev Biol. 2021;9:761709. doi:10.3389/fcell.2021.761709 PubMed PMID: 34901007; PubMed Central PMCID: PMC8652214.

96. Ibrahim L, Mohamed RH, Tolba MM, Radwan SM, Hamdy NM, Elhefnawi M. Unveiling the Therapeutic Potential of Targeting RRM2 in Hepatocellular Carcinoma: An Integrated In Silico and In Vitro Study. Funct Integr Genomics. 2025;25(1):123. doi:10.1007/s10142-025-01630-0 PubMed PMID: 40493217; PubMed Central PMCID: PMC12152083.

97. Ayoub AM, Mahmoud EAB, Mohamed RH, Tolba MM, Hassan NS, Ghazy M, et al. BIRC5 as a master regulator in HCC: unraveling its role in tumor survival and therapeutic potential. Funct Integr Genomics. 2025 Dec;25(1):120. doi:10.1007/s10142-025-01615-z

98. Schneider CA, Rasband WS, Eliceiri KW. NIH Image to ImageJ: 25 years of image analysis. Nat Methods. 2012 Jul;9(7):671–5. doi:10.1038/nmeth.2089 PubMed PMID: 22930834; PubMed Central PMCID: PMC5554542.

99. Pugazhenthi S, Nesterova A, Sable C, Heidenreich KA, Boxer LM, Heasley LE, et al. Akt/Protein Kinase B Up-regulates Bcl-2 Expression through cAMP-response Element-binding Protein. J Biol Chem. 2000 Apr;275(15):10761–6. doi:10.1074/jbc.275.15.10761

100. Mohamed NM, Mohamed RH, Kennedy JF, Elhefnawi MM, Hamdy NM. A comprehensive review and in silico analysis of the role of survivin (BIRC5) in hepatocellular carcinoma hallmarks: A step toward precision. Int J Biol Macromol. 2025 Jun;311(Pt 1):143616. doi:10.1016/j.ijbiomac.2025.143616 PubMed PMID: 40306500.

101. Guzmán C, Bagga M, Kaur A, Westermarck J, Abankwa D. ColonyArea: an ImageJ plugin to automatically quantify colony formation in clonogenic assays. PloS One. 2014;9(3):e92444. doi:10.1371/journal.pone.0092444 PubMed PMID: 24647355; PubMed Central PMCID: PMC3960247.

102. Hanahan D, Weinberg RA. Hallmarks of cancer: The next generation. Cell. 2011 Mar;144(5):646–74. doi:10.1016/j.cell.2011.02.013

103. Lakbir S, de Wit R, de Bruijn I, Kundra R, Madupuri R, Gao J, et al. Tumor break load quantitates structural variant-associated genomic instability with biological and clinical relevance across cancers. NPJ Precis Oncol. 2025 May 14;9:140. doi:10.1038/s41698-025-00922-9 PubMed PMID: 40369102; PubMed Central PMCID: PMC12078582.

104. Taddei ML, Giannoni E, Fiaschi T, Chiarugi P. Anoikis: an emerging hallmark in health and diseases. J Pathol. 2012 Jan;226(2):380–93. doi:10.1002/path.3000 PubMed PMID: 21953325.

105. Foley EA, Kapoor TM. Microtubule attachment and spindle assembly checkpoint signalling at the kinetochore. Nat Rev Mol Cell Biol. 2013 Jan;14(1):25–37. doi:10.1038/nrm3494 PubMed PMID: 23258294; PubMed Central PMCID: PMC3762224.

106. Musacchio A, Desai A. A Molecular View of Kinetochore Assembly and Function. Biology. 2017 Jan 24;6(1):5. doi:10.3390/biology6010005 PubMed PMID: 28125021; PubMed Central PMCID: PMC5371998.

107. Cheeseman IM, Hori T, Fukagawa T, Desai A. KNL1 and the CENP-H/I/K complex coordinately direct kinetochore assembly in vertebrates. Mol Biol Cell. 2008 Feb;19(2):587–94. doi:10.1091/mbc.e07-10-1051 PubMed PMID: 18045986; PubMed Central PMCID: PMC2230600.

108. Gisselsson D, Pettersson L, Höglund M, Heidenblad M, Gorunova L, Wiegant J, et al. Chromosomal breakage-fusion-bridge events cause genetic intratumor heterogeneity. Proc Natl Acad Sci U S A. 2000 May 9;97(10):5357–62. doi:10.1073/pnas.090013497 PubMed PMID: 10805796; PubMed Central PMCID: PMC25833.

109. Sansregret L, Swanton C. The Role of Aneuploidy in Cancer Evolution. Cold Spring Harb Perspect Med. 2017 Jan 3;7(1):a028373. doi:10.1101/cshperspect.a028373 PubMed PMID: 28049655; PubMed Central PMCID: PMC5204330.

110. Li J, Gao JZ, Du JL, Huang ZX, Wei LX. Increased CDC20 expression is associated with development and progression of hepatocellular carcinoma. Int J Oncol. 2014 Oct 1;45(4):1547–55. doi:10.3892/ijo.2014.2559

111. Li R, Jiang X, Zhang Y, Wang S, Chen X, Yu X, et al. Cyclin B2 Overexpression in Human Hepatocellular Carcinoma is Associated with Poor Prognosis. Arch Med Res. 2019 Jan;50(1):10–7. doi:10.1016/j.arcmed.2019.03.003 PubMed PMID: 31101236.

112. Liu S, Yuan X, Gui P, Liu R, Durojaye O, Hill DL, et al. Mad2 promotes Cyclin B2 recruitment to the kinetochore for guiding accurate mitotic checkpoint. EMBO Rep. 2022 Jun 7;23(6):e54171. doi:10.15252/embr.202154171 PubMed PMID: 35384228; PubMed Central PMCID: PMC9171689.

113. Yin S, Yang S, Luo Y, Lu J, Hu G, Wang K, et al. Cyclin-dependent kinase 1 as a potential target for lycorine against hepatocellular carcinoma. Biochem Pharmacol. 2021 Nov;193:114806. doi:10.1016/j.bcp.2021.114806

114. Yu H. Chk1: A Double Agent in Cell Cycle Checkpoints. Dev Cell. 2007 Feb 1;12(2):167–8. doi:10.1016/j.devcel.2007.01.005

115. Petsalaki E, Zachos G. Chk1 and Mps1 jointly regulate correction of merotelic kinetochore attachments. J Cell Sci. 2013 Mar 1;126(Pt 5):1235–46. doi:10.1242/jcs.119677 PubMed PMID: 23321637.

116. Li X, Huang J, Wu Q, Du Q, Wang Y, Huang Y, et al. Inhibition of Checkpoint Kinase 1 (CHK1) Upregulates Interferon Regulatory Factor 1 (IRF1) to Promote Apoptosis and Activate Anti-Tumor Immunity via MICA in Hepatocellular Carcinoma (HCC). Cancers. 2023 Jan 30;15(3):850. doi:10.3390/cancers15030850 PubMed PMID: 36765808; PubMed Central PMCID: PMC9913340.

117. Li X, Zhou M, Zhu Z, Wang Z, Zhang X, Lu L, et al. Kaempferol from Alpinia officinarum hance induces G2/M cell cycle arrest in hepatocellular carcinoma cells by regulating the ATM/CHEK2/KNL1 pathway. J Ethnopharmacol. 2024 Oct 28;333:118430. doi:10.1016/j.jep.2024.118430 PubMed PMID: 38857680.

118. Urata YN, Takeshita F, Tanaka H, Ochiya T, Takimoto M. Targeted Knockdown of the Kinetochore Protein D40/Knl-1 Inhibits Human Cancer in a p53 Status-Independent Manner. Sci Rep. 2015 Sep 8;5(1):13676. doi:10.1038/srep13676

119. Bai T, Zhao Y, Liu Y, Cai B, Dong N, Li B. Effect of KNL1 on the proliferation and apoptosis of colorectal cancer cells. Technol Cancer Res Treat. 2019 Jan 1;18:1533033819858668. doi:10.1177/1533033819858668 PubMed PMID: 31315522; PubMed Central PMCID: PMC6637841.

120. Wu L, Zhang G, Zhu Q, Huang Y. Kinetochore scaffold 1 downregulation suppressed the development of non-small cell lung cancer by inactivating the phosphatidylinositol 3 kinase/protein kinase B (AKT)/nuclear factor-kappa B pathway. J Physiol Pharmacol Off J Pol Physiol Soc. 2024 Feb;75(1). doi:10.26402/jpp.2024.1.05 PubMed PMID: 38583438.

121. Sica A, Mantovani A. Macrophage plasticity and polarization: in vivo veritas. J Clin Invest. 2012 Mar;122(3):787–95. doi:10.1172/JCI59643 PubMed PMID: 22378047; PubMed Central PMCID: PMC3287223.

122. Xue J, Schmidt SV, Sander J, Draffehn A, Krebs W, Quester I, et al. Transcriptome-based network analysis reveals a spectrum model of human macrophage activation. Immunity. 2014 Feb 20;40(2):274–88. doi:10.1016/j.immuni.2014.01.006 PubMed PMID: 24530056; PubMed Central PMCID: PMC3991396.

123. Fan Y, Chiu A, Zhao F, George JT. Understanding the interplay between extracellular matrix topology and tumor-immune interactions: Challenges and opportunities. Oncotarget. 2024 Jan 6;15:768–81. doi:10.18632/oncotarget.28666

124. Jalil SMA, Henry JC, Cameron AJM. Targets in the Tumour Matrisome to Promote Cancer Therapy Response. Cancers. 2024 Jan;16(10):10. doi:10.3390/cancers16101847

125. Wang Y, Wen X, Su C, You Y, Jiang Z, Fan Q, et al. The role of tenascin-C in tumor microenvironments and its potential as a therapeutic target. Front Cell Dev Biol. 2025 Feb 19;13. doi:10.3389/fcell.2025.1554312

126. Chen A, Shen Y, Xia M, Xu L, Pan N, Yin Y, et al. Expression of the nonclassical HLA class I and MICA/B molecules in human hepatocellular carcinoma. Neoplasma. 2011;58(5):371–6. doi:10.4149/neo_2011_05_371 PubMed PMID: 21744989.

127. Jinushi M, Takehara T, Tatsumi T, Kanto T, Miyagi T, Suzuki T, et al. Negative regulation of NK cell activities by inhibitory receptor CD94/NKG2A leads to altered NK cell-induced modulation of dendritic cell functions in chronic hepatitis C virus infection. J Immunol Baltim Md 1950. 2004 Nov 15;173(10):6072–81. doi:10.4049/jimmunol.173.10.6072 PubMed PMID: 15528343.

128. Wang XK, Liao XW, Yang CK, Yu TD, Liu ZQ, Gong YZ, et al. Diagnostic and prognostic biomarkers of Human Leukocyte Antigen complex for hepatitis B virus-related hepatocellular carcinoma. J Cancer. 2019;10(21):5173–90. doi:10.7150/jca.29655 PubMed PMID: 31602270; PubMed Central PMCID: PMC6775598.

129. Xu Y, Han H, Zhang F, Lv S, Li Z, Fang Z. Lesion human leukocyte antigen-F expression is associated with a poor prognosis in patients with hepatocellular carcinoma. Oncol Lett. 2015 Jan 1;9(1):300–4. doi:10.3892/ol.2014.2686

130. Brandt EF, Baues M, Wirtz TH, May JN, Fischer P, Beckers A, et al. Chemokine CXCL10 Modulates the Tumor Microenvironment of Fibrosis-Associated Hepatocellular Carcinoma. Int J Mol Sci. 2022 Jul 23;23(15):8112. doi:10.3390/ijms23158112 PubMed PMID: 35897689; PubMed Central PMCID: PMC9329882.

131. Halper J. Basic Components of Connective Tissues and Extracellular Matrix: Fibronectin, Fibrinogen, Laminin, Elastin, Fibrillins, Fibulins, Matrilins, Tenascins and Thrombospondins. Adv Exp Med Biol. 2021;1348:105–26. doi:10.1007/978-3-030-80614-9_4 PubMed PMID: 34807416.

132. Frisch SM, Ruoslahti E. Integrins and anoikis. Curr Opin Cell Biol. 1997 Oct;9(5):701–6. doi:10.1016/s0955-0674(97)80124-x PubMed PMID: 9330874.

133. Genda T, Sakamoto M, Ichida T, Asakura H, Hirohashi S. Loss of cell-cell contact is induced by integrin-mediated cell-substratum adhesion in highly-motile and highly-metastatic hepatocellular carcinoma cells. Lab Investig J Tech Methods Pathol. 2000 Mar;80(3):387–94. doi:10.1038/labinvest.3780043 PubMed PMID: 10744074.

134. Leng C, Zhang ZG, Chen WX, Luo HP, Song J, Dong W, et al. An integrin beta4-EGFR unit promotes hepatocellular carcinoma lung metastases by enhancing anchorage independence through activation of FAK-AKT pathway. Cancer Lett. 2016 Jun 28;376(1):188–96. doi:10.1016/j.canlet.2016.03.023 PubMed PMID: 26996299.

135. Mitra SK, Schlaepfer DD. Integrin-regulated FAK-Src signaling in normal and cancer cells. Curr Opin Cell Biol. 2006 Oct;18(5):516–23. doi:10.1016/j.ceb.2006.08.011 PubMed PMID: 16919435.

136. Wang LH. Molecular signaling regulating anchorage-independent growth of cancer cells. Mt Sinai J Med N Y. 2004 Nov;71(6):361–7. PubMed PMID: 15592654.

137. Huaman J, Naidoo M, Zang X, Ogunwobi OO. Fibronectin Regulation of Integrin B1 and SLUG in Circulating Tumor Cells. Cells. 2019 Jun 20;8(6):618. doi:10.3390/cells8060618 PubMed PMID: 31226820; PubMed Central PMCID: PMC6627780.

138. Shi Y, Shang J, Li Y, Zhong D, Zhang Z, Yang Q, et al. ITGA5 and ITGB1 contribute to Sorafenib resistance by promoting vasculogenic mimicry formation in hepatocellular carcinoma. Cancer Med. 2023;12(3):3786–96. doi:10.1002/cam4.5110

139. Kang CL, Qi B, Cai QQ, Fu LS, Yang Y, Tang C, et al. LncRNA AY promotes hepatocellular carcinoma metastasis by stimulating ITGAV transcription. Theranostics. 2019 Jun 9;9(15):4421–36. doi:10.7150/thno.32854 PubMed PMID: 31285770; PubMed Central PMCID: PMC6599657.

140. Nong Y, Wu D, Lin Y, Zhang Y, Bai L, Tang H. Tenascin-C expression is associated with poor prognosis in hepatocellular carcinoma (HCC) patients and the inflammatory cytokine TNF-α-induced TNC expression promotes migration in HCC cells. Am J Cancer Res. 2015;5(2):782–91. PubMed PMID: 25973315; PubMed Central PMCID: PMC4396033.

141. Morgan MR, Humphries MJ, Bass MD. Synergistic control of cell adhesion by integrins and syndecans. Nat Rev Mol Cell Biol. 2007 Dec;8(12):957–69. doi:10.1038/nrm2289

142. Yang H, Liu Y, Zhao MM, Guo Q, Zheng XK, Liu D, et al. Therapeutic potential of targeting membrane-spanning proteoglycan SDC4 in hepatocellular carcinoma. Cell Death Dis. 2021 May 14;12(5):492. doi:10.1038/s41419-021-03780-y

143. Yu L, Xu H, Zhang S, Chen J, Yu Z. SDC1 promotes cisplatin resistance in hepatic carcinoma cells via PI3K-AKT pathway. Hum Cell. 2020 Jul 1;33(3):721–9. doi:10.1007/s13577-020-00362-6

